# Long-read proteogenomic atlas of human neuronal differentiation reveals isoform diversity informing neurodevelopmental risk mechanisms

**DOI:** 10.64898/2025.12.16.693263

**Authors:** Nuo Xu, Katherine M. Rynard, Elizabeth Radley, Ai Tian, Maahil Arshad, Chaoying Long, Jimmy Ly, Hua Luo, Ellie Hogan, Maria Eleni Fafouti, Melanie Davie, Fatima Naimi, Yun Li, John A. Calarco, Brett Trost, Julien Muffat, Hyun O. Lee, Howard D. Lipshitz, Craig A. Smibert, Shreejoy J. Tripathy

## Abstract

RNA splicing shapes neuronal identity and disease risk, yet current maps lack the developmental resolution and depth to resolve this complexity. Here, we integrate deep long-read RNA sequencing and proteomics in iPSC-derived cortical neurons to generate a high-resolution proteogenomic atlas of human neuron development. We identify 182,371 mRNA isoforms (over half previously unknown) and provide direct peptide evidence for the translation of hundreds of novel protein-coding sequences. Population genetics demonstrates that variants affecting novel exons and splice sites are under negative selection, underscoring the potential significance of these isoforms. During neuronal maturation, we observe that ASD risk genes undergo dynamic isoform switching, including microexon inclusion and intron retention, that remodel key protein domains and regulatory regions. Furthermore, we uncover widespread, long-range coordination between splicing and polyadenylation. Finally, our atlas enables variant reinterpretation in ASD, highlighting the value of an isoform-centric view for interpreting pathogenic variation in neurodevelopment.

## Introduction

RNA splicing is an essential mechanism that generates the vast transcriptomic complexity required for human brain development, cellular identity, and function^1^. Splicing dysregulation is strongly implicated in the genetic etiology of neurodevelopmental disorders (NDDs) such as Autism Spectrum Disorder (ASD), where hundreds of risk genes are known to produce a wide array of isoforms^2–4^. However, our understanding of how isoform diversity contributes to neuronal development and disease risk remains incomplete because the full repertoire of isoforms expressed during neurodevelopment has not been resolved, fundamentally limiting our ability to interpret the impact of genetic variation^5,6^.

This knowledge gap stems from the technical limitations of short-read RNAseq, the predominant technology for transcriptomic analysis^7^. Because this method sequences only small fragments, it cannot accurately identify or quantify distinct full-length mRNA isoforms^7,8^. Consequently, the combinatorial logic linking alternative transcription start, splicing, and polyadenylation remains obscured^9^, even though targeted long-read studies indicate such features are tightly coordinated and developmentally regulated^10,11^. While recent unbiased long-read sequencing of human fetal brain tissues^12,13^ and iPSC-derived neurons^14^ have begun to catalog this diversity, these approaches lack the deep sequencing coverage required to fully resolve the precise dynamics governing these transitions.

Here, we overcome these limitations by integrating ultra-deep long-read RNAseq with mass spectrometry-based proteomics to generate a high-resolution proteogenomic atlas of human cortical neuron differentiation. Leveraging the PacBio Kinnex system^15,16^, we sequenced full-length mRNA transcripts at unprecedented read depths per sample. We developed new computational frameworks for analyzing these data, creating a foundational map of the neuronal isoform landscape that defines the dynamic regulatory principles of mRNA production during neuronal differentiation. Using this atlas, we uncover novel isoforms and provide direct proteomic and population-genetic evidence for their functional significance. Our findings demonstrate that an isoform-centric view is critical for interpreting the mechanisms of genetic risk in neurodevelopmental disorders.

## Results

### Deep characterization of human iPSC-derived neurons using multi-modal functional genomics

Human iPSC-derived neuronal models offer a powerful system that combines developmental relevance with experimental control and scalability^17^. We employed an engineered human induced pluripotent stem cell (iPSC) line from a neurotypical male donor harboring a doxycycline-inducible NGN2 construct for targeted differentiation into excitatory cortical neurons (see <u>Methods</u>). We differentiated three independent iPSC stocks (t00) into intermediate cells at 4 days in vitro (DIV; t04) and neurons at 30 DIV (t30) (Fig. 1A).

**Figure 1.**
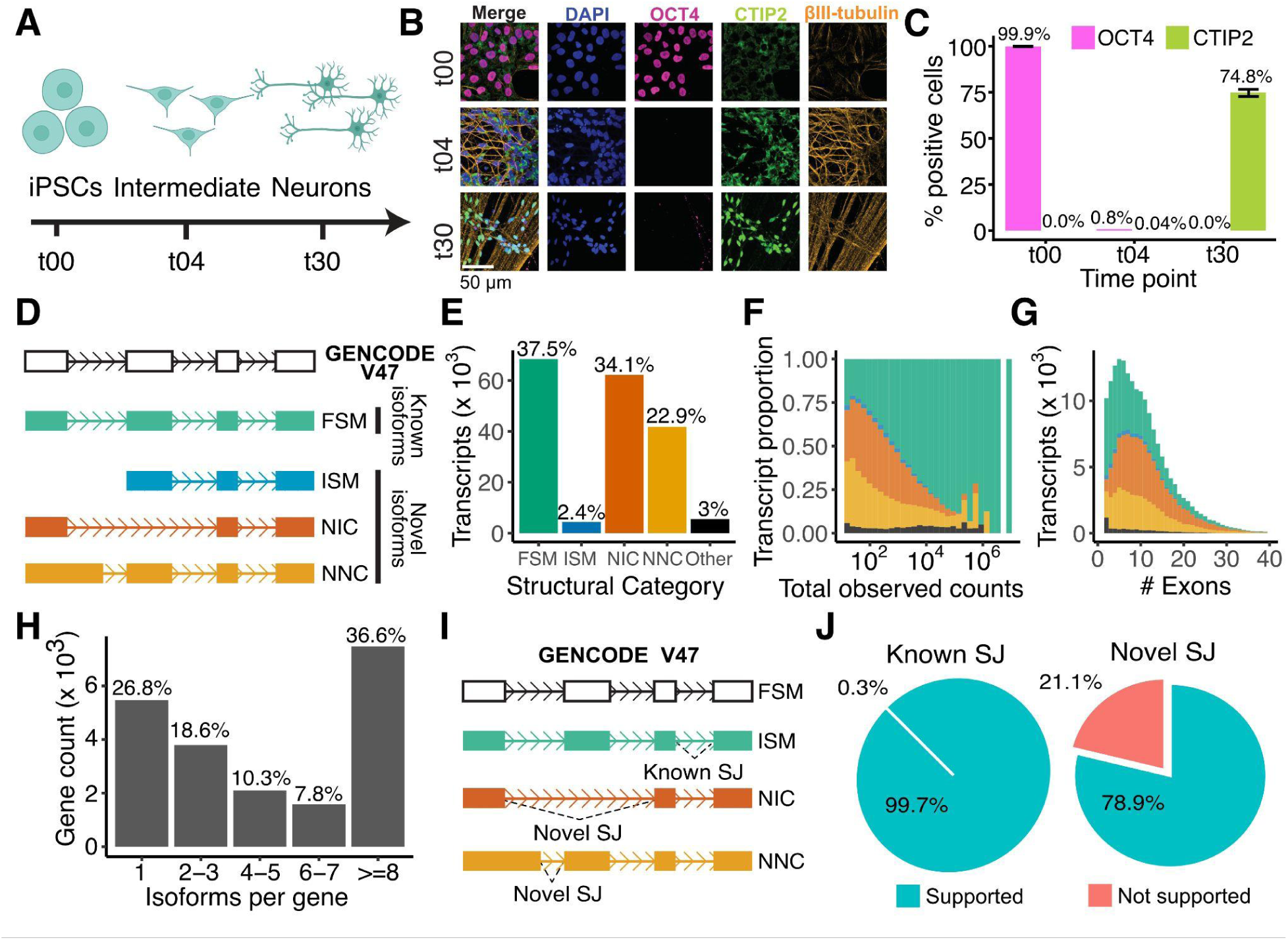
Experimental overview and landscape of expanded isoform complexity in human neuronal development. **(A)** Schematic of the experimental workflow. Human iPSCs (t00 DIV) were differentiated into an intermediate stage (t04 DIV) and then into cortical neurons (t30 DIV). Matched samples were collected at each time point for PacBio long-read RNAseq, Illumina short-read RNAseq, and shotgun proteomics. **(B, C)** Representative immunofluorescence images **(B)** and quantification **(C)** confirming cellular identity across the time course using markers for pluripotency (OCT4) and mature neurons (CTIP2). Error bars denote SEM. **(D-G)** Classification and characterization of the isoform landscape using long-read RNAseq. Isoforms are classified relative to the GENCODE V47 reference **(D)**. Categories include: Full Splice Match (FSM; matches all reference splice junctions), Incomplete Splice Match (ISM; matches consecutive but not all reference junctions), Novel In Catalog (NIC; novel combinations of known splice sites), and Novel Not in Catalog (NNC; contains at least one novel splice site). FSM isoforms are defined as ’known,’ while ISM, NIC, and NNC are defined as ’novel’. These structural categories are used to illustrate the proportion of all high-confidence isoforms **(E)**, as well as the distributions of total read counts **(F)** and exon counts **(G)** per isoform. Isoforms are colored by their structural category, as in D, throughout. **(H)** Number of unique isoforms detected per gene. **(I, J)** Definition and validation of splice junctions. The schematic in **(I)** defines splice junctions (SJs) as ’known’ if present in the GENCODE V47 reference and ’novel’ otherwise. The pie charts **(J)** show the percentage of ’known’ and ’novel’ splice junctions identified in long-read data (supported) that were independently validated by matched short-read RNAseq.

Confocal immunofluorescence confirmed cellular identity across the time course (Fig. 1B, C). From these same time points, we performed deep transcriptomic profiling using both PacBio long-read and Illumina short-read RNAseq. We further profiled the proteome using mass spectrometry and developed a novel pipeline for integrating these datasets (Fig. S1). The transcriptional signatures of our samples are consistent with those reported previously^18^, further corroborating cellular identities at each developmental stage (Fig. S2).

### Deep long-read RNAseq discovers over 100,000 novel mRNA isoforms in neuronal development

To comprehensively characterize isoform diversity, we performed ultra-deep long-read RNAseq (PacBio Revio/Kinnex) across the differentiation time course (n = 3, 6, and 6 samples from t00, t04, and t30 DIV, respectively). We generated a total of 689 million reads, representing to our knowledge one of the most deeply sequenced long-read RNAseq datasets to date. Following rigorous QC (see <u>Methods</u>, Fig. S1), we retained isoforms supported by >5 reads in ≥3 samples (median read length of ∼2.6 kb, Fig. S3)

We identified 182,371 unique mRNA isoforms across 20,442 genes (<u>Table S1</u>). We classified isoforms by comparing them to the GENCODE V47 reference^19^ using the SQANTI classification scheme^20^, which categorizes isoforms based on splice-junction agreement into four main groups: full splice match (FSM), incomplete splice match (ISM), novel in catalog (NIC), or novel not in catalog (NNC) (Fig. 1D). Because ISM transcripts can arise from technical artifacts related to RNA degradation or internal priming^20^, we retained only those with external evidence for their transcription start and end sites (see Fig. S4, <u>Methods</u>). We defined FSM transcripts as “known” and ISM, NIC, and NNC transcripts as “novel” (Fig. 1D). The isoforms can be interactively viewed and explored using our custom web app (https://nuoxuxu.shinyapps.io/transcript_vis_app/) and associated UCSC Genome Browser track (see <u>Data Availability Statement</u>).

Notably, over half of the detected transcripts (59.4%, 108,377 total) were classified as novel (Fig. 1E), revealing a vast landscape of unannotated splicing. Although novel isoforms generally showed lower expression than known ones (Fig. 1F; *P* < 2.2 x 10^-16^), they were robustly supported: 51.7% had >100 reads and 14.3% had >500 reads. Furthermore, these transcripts were highly multi-exonic (Fig. 1G). Gene-level complexity was substantial, with 38.6% of genes expressing eight or more isoforms (Fig. 1H), a two-fold increase over GENCODE V47 annotations. We also identified 1,033 microexons (<28 nucleotides) absent from GENCODE or the vastDB database^21^ (<u>Table S2</u>). Our novel isoforms showed substantial overlap (30.8%, <u>Table S1</u>) with those from an external long-read dataset from human fetal brain tissue^13^. This overlap is notable given the fundamental differences in biological context (i.e., iPSC-derived neurons versus tissue) and our greater sequencing depth, which likely captures rare isoforms missed in previous studies (Fig. S5A).

Beyond these primary splicing categories, our analysis also identified several less common transcript types (Fig. S6). A small fraction of transcripts (0.39%, or 712) originated from 632 previously unannotated “intergenic” or “antisense” gene loci (Fig. 1E-G; Other). Additionally, we identified 4,332 putative gene fusion transcripts, which are likely formed by readthrough transcription, splicing together exons from two or more distinct genes.

To validate the novel splice junctions observed, we leveraged our matched Illumina short-read RNAseq data (∼100 million 150bp paired-end reads/sample). We defined splice junctions as “novel” if they were absent the GENCODE reference and “known” if present (Fig. 1I). As expected, 99.7% of known junctions were detected in the short-read data.

Critically, 78.9% of novel junctions were independently confirmed by short reads (Fig. 1J; Fig. S5B). This high concordance between platforms provides strong evidence that the novel splicing events cataloged here represent genuine biological signals rather than technical artifacts.

### Experimental evidence confirms the translation of novel isoforms

To assess protein coding potential, we predicted open reading frames (ORFs) for the 182,371 mRNA isoforms in our long-read transcriptome using ORFanage^22^. We identified 162,913 putative coding isoforms (containing an ORF ≥50 nucleotides), which collapsed into a non-redundant set of 81,745 computationally predicted proteins (Fig. 2A). Classification using the SQANTI protein framework^23^ revealed that over half of these predicted proteins were novel (Fig. 2B,C; Fig. S7), suggesting substantial unannotated protein potential.

**Figure 2.**
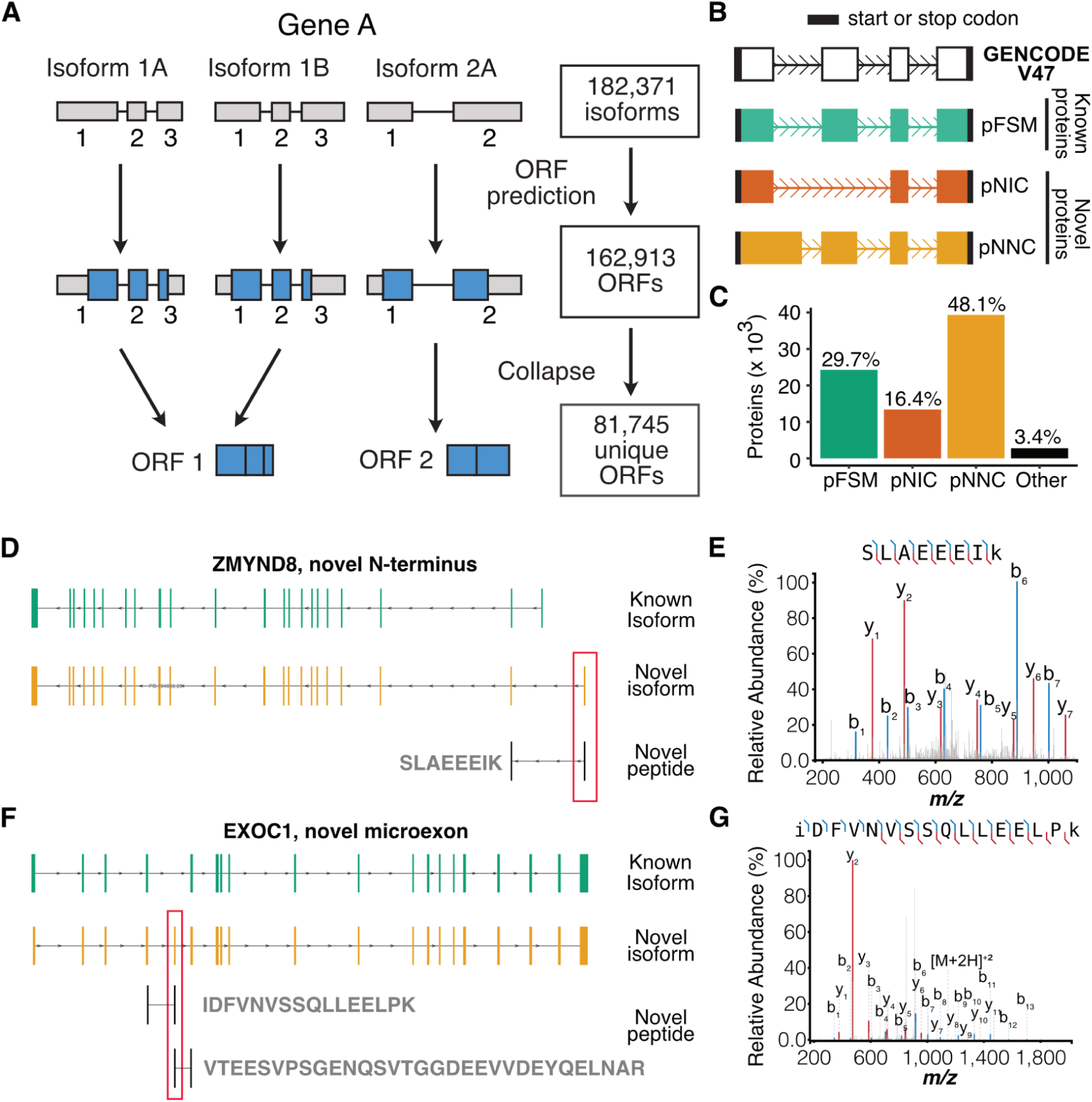
Proteogenomic validation of expanded protein isoform landscape in neurodevelopment. **(A-C)** Computational prediction and classification of protein isoforms. **(A)** Workflow illustrating the prediction of 81,745 unique ORFs from 182,371 high-confidence mRNA isoforms. **(B)** Classification of predicted protein isoforms (pFSM, pNIC, pNNC) relative to the GENCODE reference proteome. **(C)** Proportions of predicted protein isoforms across each structural category, revealing that over half are novel. **(D, E, F, G)** Mass spectrometry-based validation of predicted protein isoforms. **(D, E)** Validation of a novel translated first exon in the ASD risk gene ZMYND8. **(D)** The ZMYND8 gene locus, showing a novel isoform PB.104062.23 (orange) and the corresponding novel peptide. **(E)** Representative mass spectrum for the novel peptide SLAEEEIK, confirming its sequence and translation. Matched b- and y-ions are highlighted. (F) The EXOC1 gene locus, showing a novel isoform PB.28078.226 (orange) that contains a novel microexon that has been validated by two distinct detected peptides. (G) Representative mass spectrum for the novel peptide IDFVNVSSQLLEELPK.

We performed deep shotgun proteomics on an independent set of 11 samples across our differentiation time course to seek experimental peptide evidence supporting these predicted proteins. Given that mass spectrometry identifies proteins by measuring their constituent peptides, we used an established proteogenomic approach^23^ to search our mass spectra against a custom database integrating UniProt reference proteins^19^ with our predicted proteins defined above (see <u>Methods</u>). We defined a peptide as “novel” only if it mapped exclusively to a newly predicted protein region (i.e., by spanning a novel splice junction or originating from a novel exon).

We identified 284 distinct novel peptides, providing direct experimental evidence for the translation of 390 previously unannotated protein isoforms across 268 genes (<u>Table S3)</u>. This yield compares favorably to similar proteogenomic studies^24^ and is consistent with the observation that novel isoforms are often expressed at lower levels (Fig. S5C), limiting their proteomic recoverability. The biological significance of this approach is underscored by specific examples, such as the validation of a novel translated exon (in PB.104062.23) in the SFARI ASD risk gene *ZMYND8*, which contributes to a novel N-terminus via an alternative translation start (Fig. 2D<u>, 2E</u>). In addition, we observed strong translational evidence for a 24-nucleotide microexon in *EXOC1*, unannotated in GENCODE but present in vastDB, that was independently validated by two separate junction-spanning peptides (Fig. 2F, 2G, Fig<u>. S8</u>). Together, these data provide direct proteomic evidence that novel splice events can remodel the protein landscape of human neurons.

To further corroborate our novel ORF predictions, we re-analyzed an external Ribo-seq dataset from human iPSC-derived neurons undergoing a similar differentiation protocol^25^. As a positive control, translational efficiency (TE), defined as the ratio of ribosome protected fragments to input mRNA, was significantly greater in known GENCODE coding sequences compared to known 3’UTR regions (*P* < 2.2 x 10^-16^; Fig. S5D). Reassuringly, this pattern extended to our newly identified isoforms, where novel predicted ORFs also exhibited significantly greater TE compared to novel predicted 3’UTRs (*P* = 2.7 x 10^-9^). Together, these findings provide independent validation for the ribosomal engagement of these novel ORFs and support their translation into new proteins.

### Discordant mRNA and protein expression dynamics across neuronal differentiation

To investigate multi-omic expression dynamics during neuronal differentiation, we integrated our transcriptomic and proteomic data at the gene level. We calculated differential mRNA and protein abundance and categorized genes into nine expression trajectories based on their directional change across timepoints (defined in Fig. 3A; e.g., ‘UU’ denotes upregulation from t00-t04 and t04-t30).

**Figure 3.**
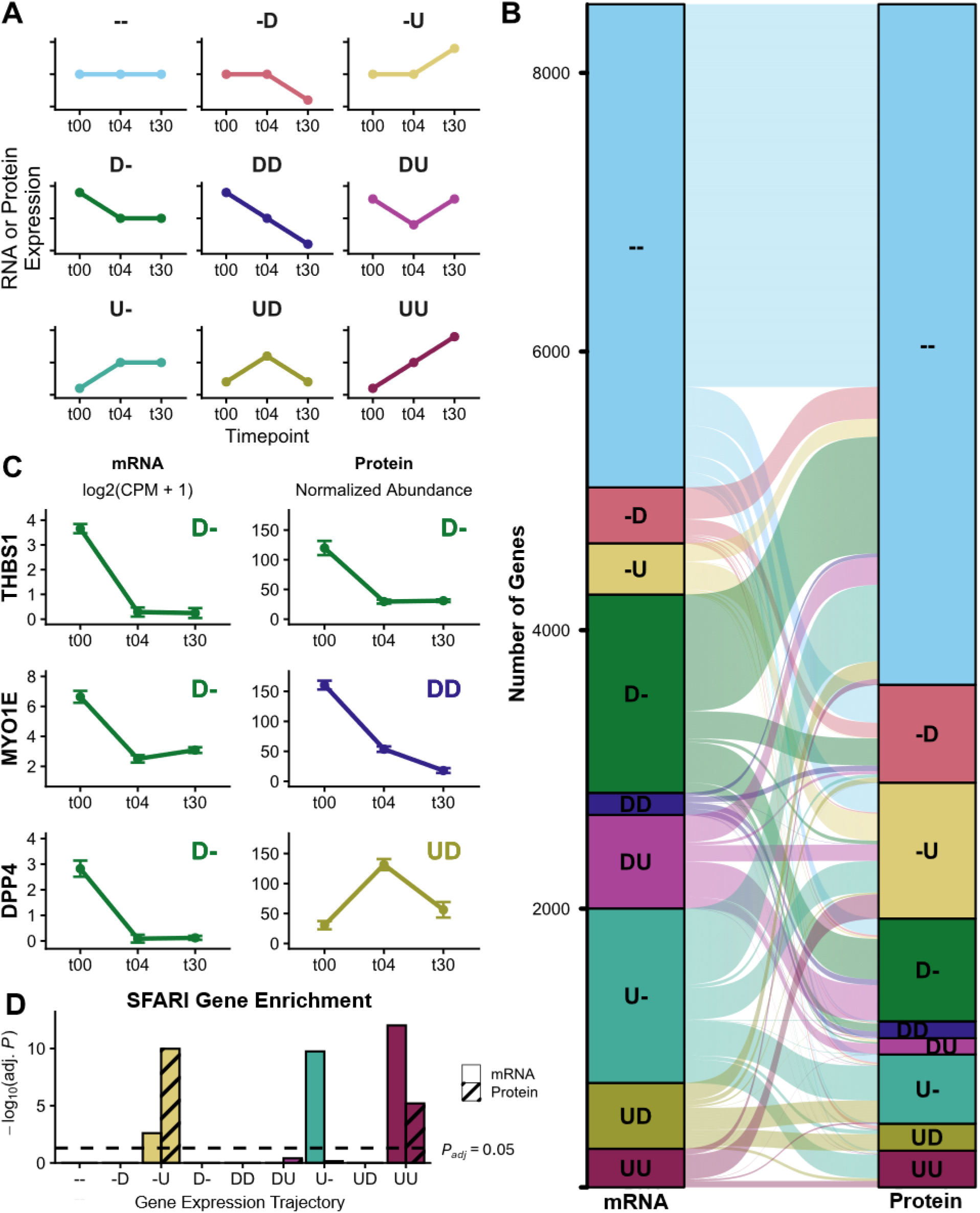
Gene-level mRNA to protein expression dynamics during neuronal differentiation. **(A)** Classification of nine expression trajectories. Trajectories are labeled with a two-character code representing the direction of change for the first (t00–t04) and second (t04–t30) intervals: ‘-’ (no change; |log2FC| < 1), ‘D’ (decrease; log2FC < -1; FDR < 0.05), and ‘U’ (increase; log2FC > 1; FDR < 0.05). **(B)** River plot mapping mRNA trajectories (left) to protein trajectories (right) for the 8,498 genes with proteomic data. **(C)** Representative examples showing concordant (THBS1), partially discordant (MYO1E), and fully discordant (DPP4) mRNA and protein dynamics. These examples are known SFARI ASD risk genes classified in the mRNA ‘D-’ trajectory. **(D)** Enrichment of SFARI ASD risk genes within mRNA and protein trajectories; dashed line indicates significance threshold (P_adj_ = 0.05).

For the 8,498 genes with both transcriptomic and proteomic data, we found extensive dynamic regulation, with 59% of mRNA transcripts and 42% of proteins changing across the time course (examples in Fig. 3B<u>, 3C</u>). While numerous genes (45.5%) showed concordant trajectories (e.g., *THBS1*), a majority exhibited discordance. Notably, 42.5% of genes differed in one timepoint comparison (e.g., *MYO1E*), and 12.0% displayed entirely discordant dynamics (e.g., *DPP4*).

Gene ontology analysis, summarized in <u>Table S4</u>, revealed that proteins with upregulated trajectories (i.e., ‘U-’, ‘-U’, ‘UU’, ‘DU’) were enriched for neuron development (GO:0048666; *P* < 9.7 x 10^-04^). Proteins that were specifically upregulated in neurons compared to intermediate cells (i.e., ‘-U’ and ‘UU’) were enriched for synapse organization (GO:0050808; *P* < 4.1 x 10^-12^). Interestingly, SFARI ASD-associated genes were enriched in the mRNA ‘U-’ trajectory, an enrichment that extended to both mRNAs and proteins in the ‘-U’ and ‘UU’ trajectories (Fig. 3D). Conversely, proteins downregulated in intermediate cells (‘D-’ trajectory) were enriched for cell cycle processes (GO:0022402; *P* = 1.1 x 10^-25^), consistent with cell cycle exit upon differentiation^26^. Overall, these high levels of discordant dynamics highlight the importance of post-transcriptional regulation. Furthermore, these analyses imply that gene-level quantification may capture only a partial view of differentiation, underscoring the need to resolve these dynamics at the isoform level.

### Widespread isoform switching reveals dynamic regulation of splicing during neuronal differentiation

To capture dynamics beyond gene-level expression, we used our long-read data to quantify changes in isoform abundance (differential transcript expression, DTE) and relative usage (differential transcript usage, DTU). We identified ∼49,000 DTE events and ∼24,000 DTU events (Fig. 4A, <u>Table S5</u>). Notably, ∼37% of the isoforms undergoing DTU were novel. We specifically focused our downstream analyses on isoform switches^27^, defined as pairwise events where the usage of one isoform increases while another decreases over time (Fig. 4B).

**Figure 4.**
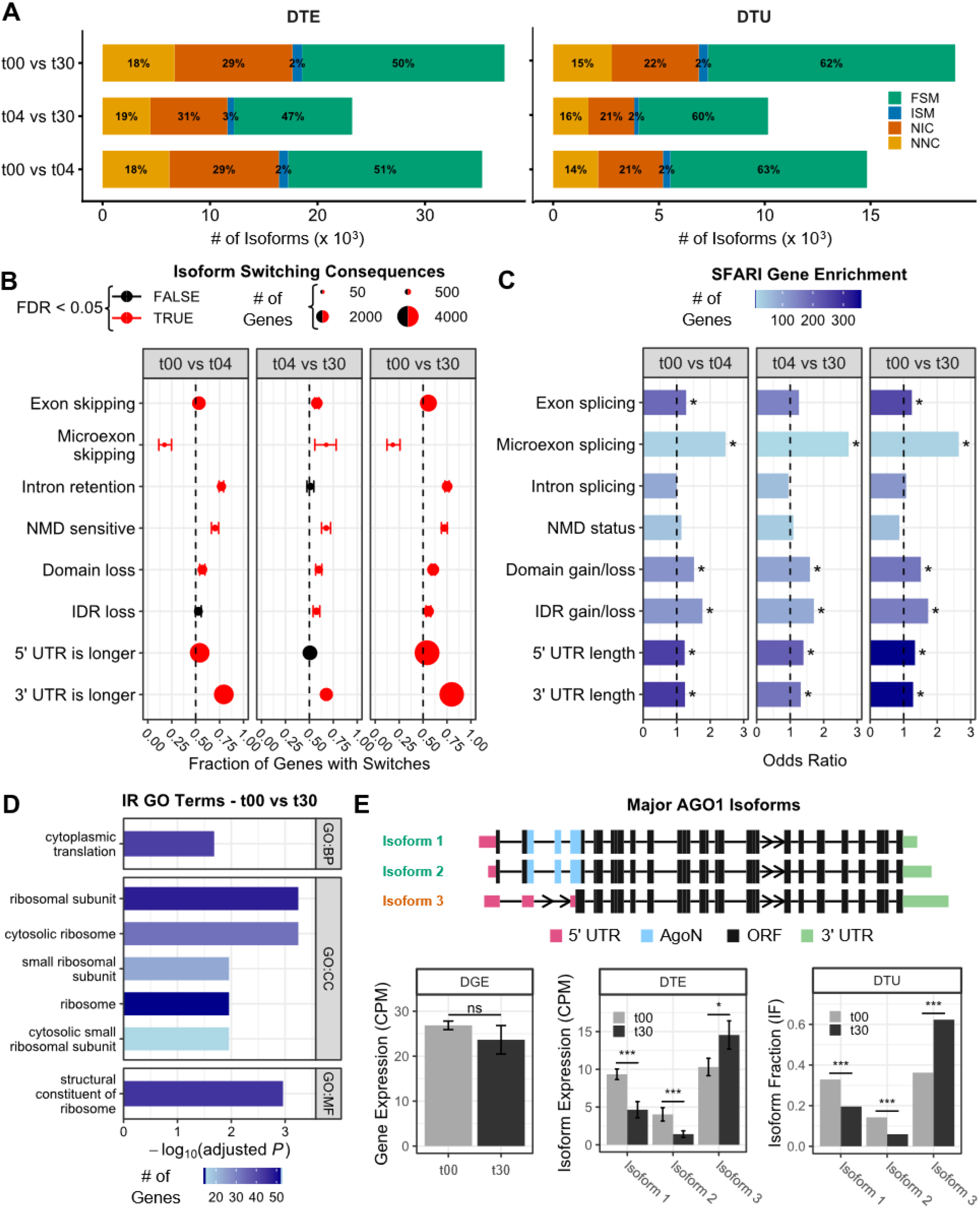
The landscape of isoform switching during cortical neuron differentiation. **(A)** Counts of differentially expressed (DTE) and differentially used (DTU) transcripts across timepoints, coloured by structural category. **(B)** Functional consequences of isoform switching. Points represent the fraction of genes favoring a specific outcome (e.g., >0.5 indicates preferential exon skipping vs. inclusion). Point size scales with the total number of genes in category. Red indicates significant bias (Binomial test, FDR < 0.05); error bars denote 95% CI. **(C)** Enrichment of SFARI ASD risk genes within genes undergoing each class of switching event in B (Fisher’s exact test; asterisk denotes FDR < 0.05). **(D)** Gene Ontology (GO) enrichment for genes regulated by intron retention (IR) from t00 to t30. **(E)** Isoform switching in the ASD risk gene AGO1. Top: Transcript models showing the switch from major known isoforms (denoted Isoform 1 and 2, green) to a novel isoform (Isoform 3, orange) which lacks the N-terminal AgoN domain. (Bottom) Quantification of DGE, DTE, and DTU for 3 major AGO1 isoforms. Error bars: 95% CI; *FDR < 0.05; ***FDR < 0.001; ns = not significant.

Systematic characterization of these isoform switches revealed distinct temporal patterns for different classes of alternative splicing (AS) events (Fig. 4C; Fig. S9; <u>Table S5</u>). We observed dynamic regulation of 179 microexons (23 of which were absent from both GENCODE V47 and vastDB). Early differentiation (t00-t04) was dominated by microexon inclusion (n=114 vs 24 genes with isoform switches resulting in inclusion vs. exclusion, FDR = 1.4 x 10^-14^). Conversely, later maturation (t04-t30) favored microexon skipping over inclusion (50 vs. 24 genes, FDR = 5.1 x 10^-03^). While previous work has emphasized microexon inclusion during differentiation^3^, our observation of late-stage microexon skipping suggests a distinct unappreciated regulatory program during neuronal maturation.

Intron retention (IR) was overwhelmingly favored early in differentiation (t00-t04; n=927 vs. 279 genes, FDR = 9.5 x 10^-81^). Because retained introns often introduce premature termination codons (PTC), we assessed nonsense-mediated decay (NMD) sensitivity. Switches resulting in PTC gain significantly increased from t00 to t30 (614 vs. 235 genes; FDR = 5.3 x 10^-39^; Fig. 4C). As expected, transcripts containing a PTC (with or without IR) showed the lower expression levels (Fig. S10). Interestingly, transcripts with IR but without a PTC displayed intermediate expression that was lower than canonical transcripts but higher than NMD targets. Genes regulated by IR were highly enriched for GO terms related to translation and ribosomal functions (Fig. 4D), consistent with the global downregulation of translation known to occur during differentiation^28^.

Isoform switches also remodeled protein structure and UTR sequences (<u>Table S5</u>). We surprisingly observed a net loss of protein domains as neurons develop (1,117 vs 729 genes with domain loss vs. gain; FDR = 5.8 x 10^-19^), particularly in metal ion binding (GO:0046872; *P* < 4.0 x 10^-04^) and dsDNA binding (GO:0003690; *P* < 0.04) genes. Similarly, switching resulted in a net loss of intrinsically disordered regions (IDRs), affecting genes involved in enzyme regulation (GO:0030234; *P* < 0.0024) and cytoskeleton protein binding (GO:0008092; *P* < 2.1 x 10^-04^). Regarding UTRs, we observed widespread 3’UTR lengthening from t00 to t30 (n=3,788 vs. 953; FDR = 3.1 x 10^-322^). In contrast, significant 5’ UTR lengthening occurred only during early differentiation (t00-t04) (Fig. 4C).

Linking these dynamics to disease genetics, we found that genes undergoing switches that altered protein domains or IDRs were significantly enriched for SFARI ASD risk genes (Fig. 4C). Notably, genes with dynamic microexon splicing showed the strongest enrichment of any category (OR = 2.6 ; *P* = 0.001 for t00-30), consistent with the established dysregulation of neuronal microexons in ASD^3^. In contrast, IR-regulated and NMD-sensitive genes showed no such enrichment.

The SFARI ASD gene *Argonaute 1* (*AGO1*), functioning in RNA-mediated gene silencing, exemplifies these multilayered dynamics. From t00 to t30, two isoform switching events result in a novel isoform comprising >60% of *AGO1* expression at t30 (Fig. 4E). These switches simultaneously lengthen both 5’ and 3’ UTRs and modify the predicted coding sequence to delete the N-terminal domain required for small RNA-AGO complex maturation^29^, the first such report of this potential protein isoform. Importantly, this remodeling was masked at the gene level, where total *AGO1* mRNA expression remained constant across timepoints, underscoring the value of isoform-level analyses.

### Alternative splicing and polyadenylation are highly coordinated during neuronal differentiation

While alternative splicing can diversify protein function^30^, alternative polyadenylation (APA) governs transcript stability, translation, and subcellular localization^31^. Long-read RNAseq uniquely enables the detection of long-range coordination between these events on single molecules (Fig. 5A). To systematically quantify such AS-APA coordination, we developed a novel statistical framework using generalized linear models (GLMs) tailored for deep, multi-sample datasets (see <u>Methods</u>). In contrast to contingency table-based approaches^10,32^, our approach models sample-to-sample variance, enabling robust, transcriptome-wide detection of dynamic AS-APA coordination.

**Figure 5.**
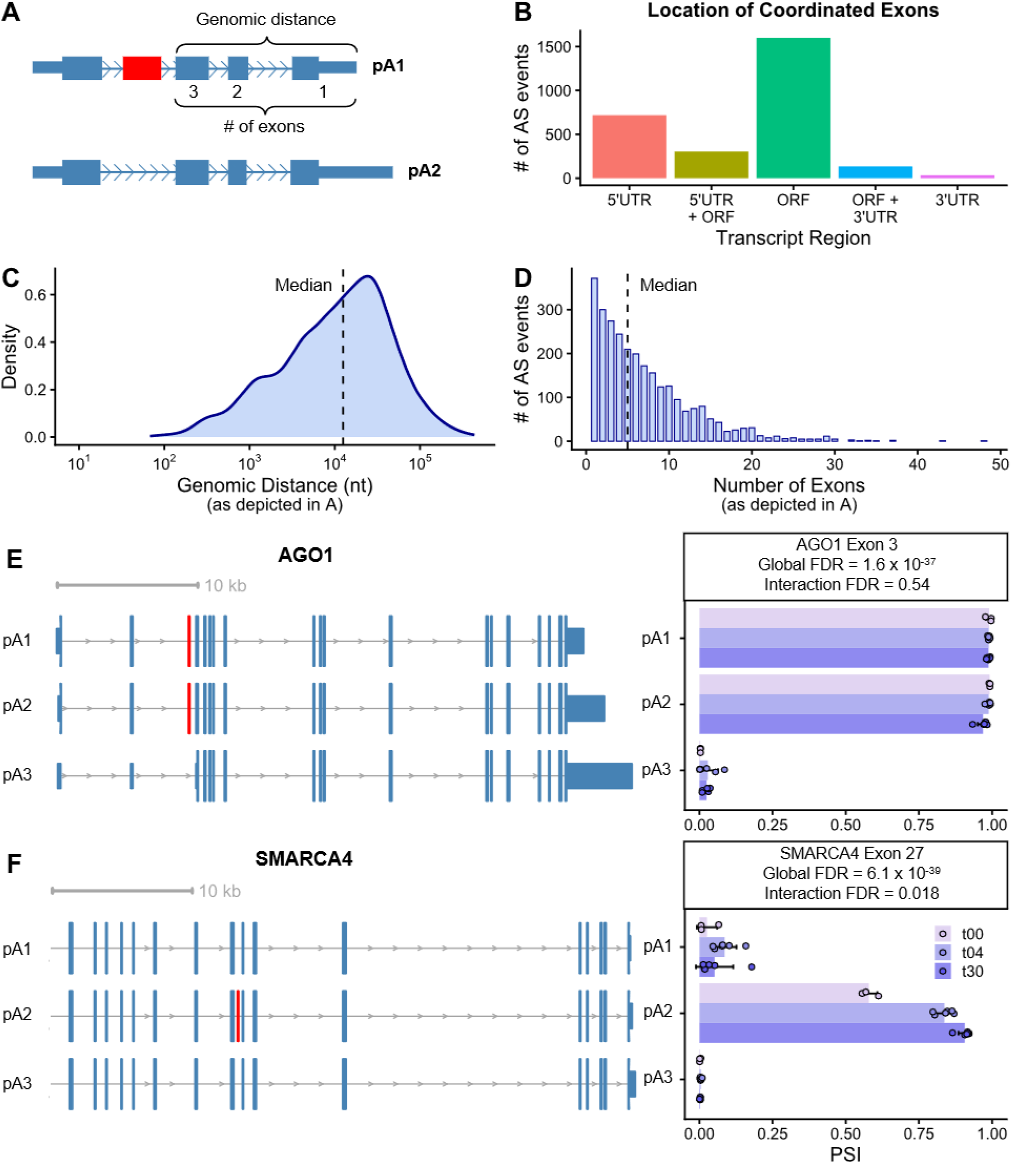
Long-range coordination of alternative splicing and polyadenylation during neuronal differentiation. **(A)** Schematic of an AS-APA coordination event. This analysis utilizes full-length reads to quantify the statistical dependency between the inclusion of an alternatively spliced (AS) exon (red) and the usage of specific downstream alternative polyadenylation (APA) sites (pA1 vs. pA2) within the same transcript. Brackets indicate the criteria for measuring genomic distance (**C**) and number of intervening exons (**D**). **(B)** Genomic context of coordinated AS exons, annotated based on the most abundant isoform. **(C, D)** Distributions of genomic distances (**C**) and number of intervening exons (**D**) between the AS exon and the most proximal polyadenylation (polyA) site. **(E, F)** Representative AS-APA coordination events in the ASD risk genes, AGO1 (**E**) and SMARCA4 (**F**). (Left) Representative transcript tracks for the three most abundant polyA sites (pA1, proximal to pA3, distal); the coordinated AS exon is highlighted in red. Tracks for SMARCA4 display the 3’ end of the gene for visual clarity. (Right) Sample-specific Percent Spliced In (PSI) values for the AS exon across polyA sites and timepoints. Error bars denote standard deviations. Inset FDR values indicate significance for global (stable) and interaction (time-dependent) coordination models.

We assessed two coordination types: “global” effects, i.e., a stable association between polyA (pA) site and exon inclusion; and “interaction” effects, where such coordination changes across time. We identified 2,789 significant AS-APA coordination events (FDR < 0.05 and ΔPSI ≥ 0.1; PSI = percent spliced in; <u>Table S6</u>). These events were in 1,606 genes (of 3,392 genes identified with the potential for such regulation) and were significantly enriched for SFARI ASD risk genes (*P* = 0.01). Most coordinated exons were located within the ORF (Fig. 5B), indicating that this mechanism simultaneously regulates protein-coding and 3’UTR sequences. Coordination often spanned large genomic distances (median 12,527 nucleotides; Fig. 5C) and multiple intervening exons (median of 5 exons, and even up to 48 exons; Fig. 5D).

The ASD risk gene, *AGO1*, exemplified a significant ‘global’ coordination event (FDR = 1.6 x 10^-37^; Fig. 5E), where exon 3 was nearly constitutively included when proximal pA sites were used (∼98% PSI) but was almost always skipped when the most distal pA site was used (∼2.2% PSI). In contrast, *SMARCA4*, another ASD risk gene, exhibited a significant interaction event (FDR = 0.018; Fig. 5F). Exon 27 inclusion was negligible with pA1 and pA3 usage but was regulated in a time-dependent manner when paired with pA2, increasing from 58 to 91% PSI from t00 to t30. Finally, comparisons between our novel method for assessing AS-APA coordination against more traditional contingency-table approaches yielded a 65% overlap between methods (<u>Table S6</u>; see <u>Methods</u>), demonstrating robustness of these findings.

Given the prevalence of 3’UTR lengthening as neurons develop (Fig 4B), we tested if these changes correlated with mRNA abundance, a relationship that has been debated in the literature^31,33^. We assessed this at both the whole-gene level and within transcript groups sharing identical coding sequences to isolate 3’UTR effects. We found no significant correlation in either analysis (Fig. S11; <u>Table S5</u>). This finding aligns with recent studies in neurons^34,35^, suggesting that UTR lengthening in this context is not globally biased towards RNA stabilization or destabilization.

### Novel exons and splice sites are mutationally constrained and contribute to ASD risk

Finally, we hypothesized that the novel coding regions and splice sites discovered in our atlas may represent functionally constrained elements whose disruption contributes to disease. To test this, we utilized population genetics to quantify their mutational constraint and assessed the impact of *de novo* mutations within them on ASD risk.

First, to determine if variants disrupting our novel predicted protein-coding regions are under negative or purifying selection, we calculated Mutability-Adjusted Proportion of Singletons (MAPS) scores^36^ using genetic variation from 730,947 individuals in the gnomAD v4.1 database with whole-exome sequencing (WES)^37,38^. We restricted this analysis to regions lacking any protein annotation in GENCODE, enabling us to test for constraint in areas traditionally classified as non-coding. As a positive control, variants within known protein-coding regions showed the expected strong gradient of constraint (nonsense > missense > synonymous) (Fig. 6A). Crucially, although the absolute magnitude of constraint was lower than in known coding sequences, our novel regions mirrored this overall pattern, where nonsense (MAPS = 0.08, FDR = 2.89 x 10⁻^4^) and missense (MAPS = 0.024, FDR = 3.69 x 10⁻^9^) variants within newly identified ORFs were significantly more mutationally constrained than synonymous variants (Fig. 6A). We next examined essential splice sites (the two intronic bases flanking exons); here, we defined splice site novelty strictly by the absence of the splice junction in the reference annotation (as in Fig. 1), meaning that we retained splice sites even if they resided within known protein-coding exons. While constraint at known essential splice sites was exceptionally strong as expected^36^, our novel essential splice sites also showed a robust signal of constraint. We confirmed these overall patterns using gnomAD whole-genome sequencing (WGS) data (Fig. S12). Together, these data show that the genomic sequences encoding our novel transcripts are under significant negative selection, supporting their biological relevance within the human population.

**Figure 6.**
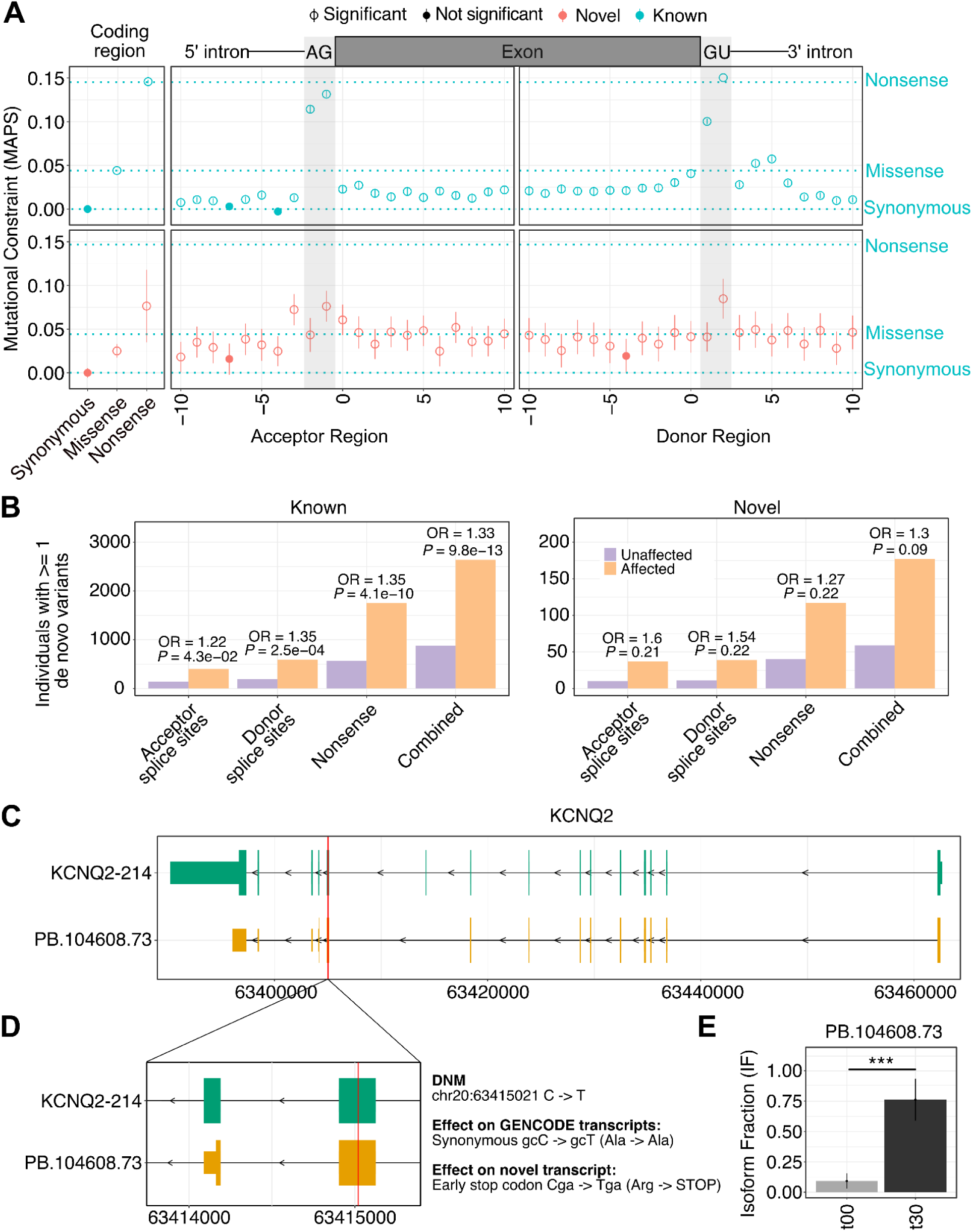
Novel exons and splice sites are functionally constrained and contribute to ASD risk. **(A)** Mutational constraint at known (top) and novel (bottom) coding sequences and near splice sites. Plots display mutational constraint (MAPS scores) for single nucleotide variants. (Left) Coding consequences for synonymous, missense, and nonsense variants. (Right) Splice regions; shaded areas highlight the essential splice sites. Open circles indicate significant constraint relative to synonymous variants (FDR < 0.05, chi-squared test). Blue dashed horizontal lines indicate MAPS scores from known elements (as in top left) shown for context. Error bars represent 95% CIs. **(B)** Burden of disruptive de novo mutations in ASD individuals versus unaffected siblings for known (left) and novel (right) elements. Odds ratios (OR) and P-values are shown (Fisher’s exact test). **(C-D)** Variant reclassification in the ASD risk gene KCNQ2. A de novo mutation (red line), classified as synonymous in the known isoform (green, KCNQ2-214), is reclassified as nonsense in a novel isoform (orange, PB.104608.73) within the alternative reading frame resulting from exon skipping. **(E)** Significant upregulation of the novel KCNQ2 isoform during neuronal maturation (***P < 0.001).

Building on this evidence of constraint, we next tested if *de novo* mutations (DNMs) reclassified as disruptive (nonsense and essential splice site) by our expanded transcriptome atlas are enriched in ASD. We analyzed a combined cohort of 94,649 individuals from the MSSNG, SSC, and SPARK studies^39–41^, comparing individuals with ASD to unaffected siblings (∼94% analyzed by WES). As a positive control, DNMs disrupting known genomic elements were highly enriched in individuals with ASD (Odds Ratio, OR = 1.33, *P* = 9.8×10⁻^13^; Fig. 6B). We then assessed DNMs considered lower-impact by GENCODE (e.g., intronic or missense) but reclassified as disruptive solely based on our novel expanded transcriptome atlas. This reclassified set displayed a comparable effect size (OR = 1.30; Fig. 6B) and a trend toward enrichment (*P* = 0.09; Fig. S13; DNM variants in <u>Table S7</u>). While this enrichment did not reach statistical significance, likely due to the rarity of such reclassified variants, the similar overall odds ratio to that of known variants suggests that DNMs impacting these novel elements contribute to ASD risk.

Crucially, our expanded isoform catalog provides a reference for reinterpreting individual risk variants. For instance, a DNM in the ASD risk gene *KCNQ2* is classified as a synonymous variant based on GENCODE (Fig. 6D). However, our data reveal a novel *KCNQ2* isoform (PB.104608.73) where this same variant introduces a nonsense codon (Arg -> STOP) within the altered reading frame resulting from exon skipping. Notably, this novel isoform is significantly upregulated during neuronal maturation (Fig. 6E), underscoring its potential functional relevance. These results demonstrate that our expanded isoform catalog can help reclassify previously overlooked mutations to uncover novel genetic risk mechanisms in ASD.

## Discussion

Here, we present the most deeply sequenced isoform-resolved map of human cortical neuron differentiation to date, integrating long-read transcriptomics with proteomics and short-read validation. Our work uncovers a vast landscape of over 100,000 novel mRNA isoforms, demonstrating that the complexity of the neuronal transcriptome, long recognized as the most complex of any tissue^42^, has nevertheless been substantially underestimated. We provide robust evidence that a significant subset of these novel isoforms is not transcriptional noise but is likely functionally significant: these isoforms are translated into protein, are dynamically regulated during differentiation, exhibit significant mutational constraint, and harbor genetic variants that may contribute to ASD risk.

A central contribution of this work is the rigorous validation of novel protein-coding isoforms. By coupling our expanded transcriptome with deep proteomics, we provided direct peptide evidence for the translation of 390 novel protein isoforms. Furthermore, our population genetics analysis is the first to apply mutational constraint metrics to a large set of novel predicted coding regions and splice sites, revealing that they are under significant negative selection. The concordance of our constraint metrics for known splice sites with previous large-scale studies provides strong validation for our approach^36^, which builds on foundational concepts of constraint from population-scale sequencing^43^. Together, these orthogonal lines of evidence demonstrate that many of the novel isoforms identified here likely represent bona fide, functionally relevant protein-coding transcripts.

That said, a substantial fraction of the novel isoforms we identified lack strong multi-omic or cross-dataset support, particularly those expressed at very low levels (Fig. S5). The biological nature of this long tail of low-level transcripts has been hotly debated^44^. A considerable portion of these molecules may represent transcriptional noise combined with the byproducts of complex and imperfect splicing, cleavage, and polyadenylation machineries^45,46^. However, this group may also contain transcripts that are more abundant and functionally relevant in specific cell types, developmental phases, and/or in response to particular environmental cues. Distinguishing among these possibilities remains a key challenge for the field.

Our time course analysis reveals that isoform switching is a pervasive regulatory mechanism driving novel splicing dynamics as neurons develop. First, while microexons are strongly linked to ASD and are generally thought to be included as neurons differentiate^3^, here we have identified that microexons exhibit a distinct biphasic pattern: inclusion during early phases followed by skipping during later maturation. This suggests the existence of a previously unappreciated, stage-specific microexon regulatory program. Second, we observed widespread intron retention linked to the downregulation of ribosomal protein genes. This provides a potential mechanism for tuning translational capacity, which may act in conjunction with or independent of the documented role of mTOR in downregulating protein synthesis during differentiation^28^. Third, isoform switches frequently remodeled predicted protein domains, particularly in ion- and DNA-binding genes, consistent with splicing’s known role in diversifying ion channels and transcription factors^47,48^. Finally, our deep long-read sequencing uncovered extensive coordination between alternative splicing and polyadenylation, often linking events separated by thousands of nucleotides. Unraveling the precise combinatorial logic of how RNA-binding proteins orchestrate these diverse regulatory programs represents an exciting frontier for future mechanistic studies.

The clinical relevance of this expanded isoform catalog is underscored by its ability to refine our understanding of ASD genetics. Our analyses demonstrate that genes regulated by specific isoform switching events, particularly those involving microexons and protein domains, are enriched for known ASD risk genes. Crucially, our atlas provides a framework for reinterpreting genetic variation in neurodevelopmental disorders. By annotating variants against our comprehensive isoform models, we show that specific mutations previously classified as non-coding or benign can be reclassified as high-impact, disrupting novel predicted protein-coding exons and essential splice sites. This reclassification provides a mechanism to resolve previously unexplained cases of ASD. Our approach extends the paradigm of using transcriptome sequencing to solve Mendelian diseases^5,49,50^ and provides a resource to identify novel pathogenic mechanisms, such as those driven by deep intronic variants^51^, effectively expanding the functional exome beyond current reference annotations.

### Limitations of the Study

We acknowledge several limitations. First, our findings rely on an iPSC-derived model of cortical neuron development. While this system offers precise molecular control, it does not capture the full cellular diversity, 3D architecture, or complex signaling environment of the *in vivo* human brain. Second, our atlas was generated from a single male donor; further studies across diverse genetic backgrounds are necessary to establish the generalizability of these specific isoforms^52^. Third, many of our functional annotations, such as ORF and protein domain predictions, are computational. For example, while we predict a functional domain loss in the critical ASD risk gene *AGO1*, future direct experimental validation of the novel start codon usage will be required. Finally, while our proteomics validation was extensive, it still represents only a fraction of the predicted novel proteome.

## Acknowledgements

This work was supported by a grant from the Simons Foundation Autism Research Initiative (SFARI #877745, H.D.L.). N.X., S.J.T., E.H., M.E.F., M.D. additionally acknowledge the CAMH Discovery Fund, Krembil Foundation, Natural Sciences and Engineering Research Council of Canada (RGPIN-2020-05834 and DGECR-2020-00048), and Canadian Institutes of Health Research (PJT-191747, NGN-171423, and PJT-175254), and Brain Canada (Future Leaders in Canadian Brain Research grant to S.J.T.). E.R. and H.O.L acknowledge funding from CIHR PJT-175174 and Canada Research Chairs program. This work was additionally supported by a grant from the Canadian Institutes of Health Research, PJT-159494 (to C.A.S.). K.M.R. was partially supported by a University of Toronto Open Fellowship. J.L. is supported in part by NSERC-PGSD and NIGMS grants (R35GM126930). For this work, J.M. acknowledges funding from the Canada Research Chair program, the Natural Sciences and Engineering Research Council of Canada (RGPIN-2019-06938), the Stem Cell Network (ECR-C4R1-4) and Brain Canada/the Azrieli Foundation. A.T. is a recipient of a 2024 NARSAD/BBR Young Investigator award.

We thank Elizabeth Tseng and Andrew Mason from PacBio for useful discussions and Shamsuddin Bhuiyan and Paul Pavlidis for feedback on the manuscript. We acknowledge the use of large language models, including Google’s Gemini Pro, in the drafting and editing of this manuscript.

## Data / Code Availability Statement

Mass spectrometry data are available via ProteomeXchange with identifier PXD069736. Raw long-read and short-read RNAseq data have been deposited to Gene Expression Omnibus with identifiers GSE311240 and GSE310131. All code used to process the data and generate the figures is available at https://github.com/nuoxuxu/SFARI. Processed data files are available at https://zenodo.org/records/16897791. A web application for viewing the full-length isoforms and their abundances across time points is available at https://nuoxuxu.shinyapps.io/transcript_vis_app/. A UCSC genome browser track hub that contains the full-length isoforms and peptides detected from proteomics is available at https://genome.ucsc.edu/s/nuoxuxu/iPSC_Neuron_Proteogenomic_atlas.

## Author Contributions

H.D.L., C.A.S., S.J.T., J.A.C., H.O.L., and J.M. jointly conceived the study and designed the experiments. A.T., C.L., and F.N. generated the iPSC lines and developed the culture conditions and differentiation protocols. E.R. scaled up, differentiated, characterized, and collected iPSC and iPSC-derived neurons and prepared samples for mass spectrometry. H.L. isolated RNA for RNAseq. N.X. and K.M.R. carried out the bulk of the data analysis and the writing of the first draft including figures, with the assistance of E.R. M.A. led the ASD burden analysis. J.L. and E.H. carried out Ribo-seq analyses. M.D., E.H. and M.E.F. contributed to the analyses of short-read RNAseq data. S.J.T., C.A.S., and H.D.L. coordinated the research, analytical design/implementation, and the writing of the manuscript. Supervision: N.X., E.H. and M.E.F by S.J.T.; K.M.R. by C.A.S.; E.R. by H.O.L; A.T., C.L and F.N. by J.M. and Y.L.; H.L. by H.D.L.; M.A. by B.T. All authors reviewed, edited, and approved the final version of the manuscript.

## Methods

### NGN2-inducible iPSC line generation

PGPC17_11 iPSCs^53^ were used as the parental iPSC line for genetic engineering. The CLYBL safe harbor sgRNA (ATGTTGGAAGGATGAGGAAA, R. Tian et al., 2019) was cloned into the pX330-GFP backbone^55,56^. iPSCs were dissociated into single cells and electroporated with both pCLYBL-TO-hNGN2-BSD-mApple (Addgene #124229, *Tian et al., 2019*) and pX330-GFP-CLYBLsgRNA. Electroporated iPSCs were selected with Blasticidin (Wisent, 400-190-EM) for two days. Integration of TO-hNGN2-BSD-mApple was further assessed by persistent mApple fluorescence. Individual mApple-positive iPSC colonies plated at clonal densities were manually picked and expanded. We verified mApple insertion and correct targeting of TO-hNGN2 into the CLYBL locus in candidate clones by PCR. A clone with successful integration of TO-hNGN2-BSD-mApple into the CLYBL locus was selected for this work.

#### iPSC culture

PGPC17_11-CLYBL-TetO-NGN2 iPSCs were grown in 6-well plates coated with Matrigel (#354234, Corning) in mTeSR Plus media (#100-0276, STEMCELL) and passaged with ReLeSR (#100-0483, STEMCELL) for routine maintenance or Accutase (#A1110501, Thermo Fisher Scientific) for single-cell plating. All cells were kept at 37°C, 95% humidity and 5% CO_2_. Matrigel coating was conducted overnight at room temperature then stored at 4°C until use according to the manufacturer’s instructions. For immunofluorescence, cells were grown on 12 mm glass coverslips coated with polyethyleneimine (PEI; #P3143, Millipore Sigma). Coverslips were first washed with 70% EtOH for 30 minutes inside a BSC then washed twice with TC ddH2O. Glass coverslips were coated with twice the recommended concentration of Matrigel. The surface was covered with sterile filtered 0.1% PEI syrup and incubated overnight at 4°C. The PEI was washed off with sterile PBS then five times with TC ddH_2_O and then dried.

#### Neuron differentiation and culture

iPSCs were dissociated from maintenance culture plates with Accutase and seeded on Matrigel-coated 6-well plates or glass coverslips at a density of 260,000 cells/cm^2^ in custom complete neuron media with 10 μM ROCK inhibitor (Y-27632, #AB144494, Abcam) and 2 μg/mL doxycycline hyclate (DOX, #D9891, Sigma Aldrich) to induce NGN2 over-expression and neuron differentiation. ROCK inhibitor was removed after 24 hours. Media was changed every 24 hours with fresh complete neuron media with DOX. On t04 cells were passaged by Accutase and plated at 260,000 cells/cm^2^ on PEI-coated (#P3143, Millipore Sigma) 6-well plates or glassware as needed in complete neuron medium with ROCK inhibitor and DOX. ROCK inhibitor was removed after 24 hours. Media was changed every 24 hours with fresh complete neuron media with DOX until t07, after this, DOX was removed, and media was changed daily until t30. Neuron media was prepared as a 25 mL custom supplement added to 500 mL Neurocell media with 3 g/L NaCl (#305-015-CL, Wisent) and stored at 4°C for a maximum of two weeks as the complete medium. The supplement consisted of 10x W21 without vitamin A (#003-014-XL, Wisent), 10x N2 (#305-016-IL, Wisent), 140 mg/mL biotin (#B4639, Millipore Sigma), 60 μg/mL NaCl (#S5886, Millipore Sigma), 100 μg/mL L-ascorbic acid 2-phosphate sesquimagnesium salt hydrate (#A8960, Sigma Aldrich), 40 μg/mL AlbuMAX-1 (#11020021, Thermo Fisher Scientific), 20 mM sodium pyruvate (#11360070, Gibco), 40 mM Gluta-Plus (609-066-EL, Wisent), 3.6% v/v NaOH 1.0N (#S2770, Sigma Aldrich), 0.4% (v/v) DL-lactic acid syrup (#L1250, Sigma Aldrich). The supplement was sterilised by double filtration and frozen at -20°C until use. All cells were kept at 37°C, 95% humidity and 5% CO_2_. Three independent iPSC cultures derived from a single donor clone were each differentiated to t04 and t30.

#### Immunofluorescence and microscopy

Cells grown on glass coverslips were imaged by confocal immunofluorescence on t00 and t30. Coverslips were PBS washed, fixed with 4% paraformaldehyde, permeabilised with 0.3% Triton X-100 and blocked with 3% BSA 0.05% Tween-20, all in PBS. Primary antibody in blocking solution was incubated at 4°C overnight, and secondary antibody in blocking solution was incubated for 1 hour at room temperature away from light. Coverslips were DAPI stained in methanol at room temperature away from light, mounted and sealed before storage at 4°C in the dark until imaging. Images were acquired on a Nikon Ti-2E microscope with an Andor Dragonfly spinning disk using a 60x oil immersion objective and a Zyla sCMOS camera after excitation with 405nm, 521 nm, 600 nm, and/or 700 nm lasers.

Primary antibodies: 0.3 μg/mL chicken anti-βIII tubulin (#AB9354, Sigma Aldrich), 1 μg/mL rabbit anti-*OCT4* (#ab19857, Abcam), 1 μg/mL mouse anti-*CTIP2* (BCL11B) (#MABE882, Sigma Aldrich). Secondary antibodies: goat anti-Mouse IgG Alexa Fluor^TM^-488 (#A11029), goat anti-Chicken IgG Alexa Fluor^TM^-555 (#A21437), goat anti-Rabbit IgG Alexa Fluor^TM^-647 (#A21245). All secondary antibodies were from Thermo Fisher and used at 2 μg/mL.

Three images from each replicate were analysed in ImageJ to confirm neuronal differentiation and determine population enrichment for cortical excitatory lineage neurons. DAPI-positive nuclei were manually marked as ROI in ImageJ. The same ROI were superimposed over the *OCT4* and *CTIP2* channels, and nuclei that were positive for either were counted. As *CTIP2* has been observed in the cell bodies of immature or non-cortical neuron cells^57,58^, any cell that showed significant *CTIP2* signal outside of the ROI was discounted. From this the percentage of *OCT4* or *CTIP2* positive-nuclei was calculated per image, averaged for each replicate and standard deviation calculated and Welch’s t-test applied.

#### RNAseq sample preparation

At day 0 (iPSC, t00), day 4 (intermediate, t04), and day 30 (neuron, t30), cells were dissociated into a single-cell suspension using Accutase (#A1110501, Thermo Fisher Scientific). Following dissociation, cells were pelleted by centrifugation, washed with sterile PBS, and resuspended at a concentration of 10⁶ cells in 0.5 mL of TRIzol™ Reagent (#15596018, Thermo Fisher Scientific). Samples were then stored at -80°C until processing.

The experimental design was initiated from three iPSC stocks, which served as the n=3 samples for the t00 timepoint. These stocks were differentiated at staggered times, with each of the three differentiations plated into triplicate wells. For the t04 and t30 timepoints, the two wells from each iPSC stock with the highest RNA concentrations were selected for sequencing library preparation. This resulted in three iPSC samples, six intermediate samples, and six neuron samples.

For long-read sequencing, full-length cDNA was first synthesized and amplified from total RNA using the Iso-Seq Express 2.0 Kit (PacBio). The resulting full-length cDNA was then used to prepare SMRTbell libraries with the Kinnex Full-Length RNA Kit (PacBio). Each library was sequenced on a single 25M SMRT Cell on the PacBio Revio sequencing system.

For short-read RNAseq, sequencing libraries were prepared using the stranded rRNA-depletion workflow (Ribo-Zero Plus rRNA Kit; Illumina), which removes bacterial and mammalian rRNA as well as human globin transcripts. Sequencing was performed on an Illumina NovaSeq 6000 platform using an S4 flowcell to generate paired-end 150 bp reads, with a target depth of approximately 100 million reads per sample.

#### Proteomics via TMT-MS

For proteomic analysis, cells at each differentiation timepoint were washed with ice-cold sterile PBS and dissociated directly in 6-well plates by incubation with lysis buffer (200 mM EPPS pH 8.5, 8 M urea, 0.25% SDS) supplemented with 1x protease inhibitor cocktail (Roche, #11836170001). To shear genomic DNA, the resulting lysate was passed through a 21-gauge needle. Total protein concentration was determined using the Pierce BCA Protein Assay Kit (Thermo Fisher Scientific, #23225) according to the manufacturer’s instructions. Samples were normalized to a minimum of 1 mg/mL, with a total protein amount of at least 200 μg prepared for each. Final aliquots were stored at -80°C until they were shipped for full proteome profiling.

Mass spectrometry analysis was performed by the Thermo Fisher Center for Multiplexed Proteomics (TCMP) at Harvard Medical School. From each submitted sample, 50 μg of protein was processed. Proteins were reduced with tris(2-carboxyethyl)phosphine (TCEP), alkylated with iodoacetamide, and the reaction was subsequently quenched with dithiothreitol (DTT). The proteins were then precipitated onto SP3 beads for buffer exchange and cleanup prior to enzymatic digestion. Digestion was carried out sequentially, first with Lys-C overnight, followed by a 6-hour incubation with trypsin. The resulting peptides were conjugated with 11-plex isobaric tandem mass tags (TMT), pooled into a single multiplexed sample, and desalted. The pooled peptides were fractionated into 24 fractions using basic reverse-phase high-performance liquid chromatography (HPLC).

Twelve of these 24 fractions were analyzed by TMT-MS on an Orbitrap Lumos Mass Spectrometer (Thermo Fisher Scientific). Data were acquired using a data-dependent acquisition (DDA) method selecting the top 10 most abundant precursor ions. MS2 scans were performed at a resolution of 50,000 with an automatic gain control (AGC) target of 2e5 and a maximum injection time of 86 ms. MS3 scans for reporter ion quantification were performed at a resolution of 50,000 with an AGC target of 1.2e5 and a maximum injection time of 105 ms. A normalized collision energy (NCE) of 36% was used for higher-energy collisional dissociation (HCD) fragmentation. Peptide identification and protein quantification were performed using the TCMP’s standard data analysis pipeline (see details below).

### Functional Genomics Data Analysis

#### Long-read RNAseq data processing

Kinnex data were preprocessed on-instrument on a PacBio Revio. Basecalling used the PacBio basecaller (default Revio parameters); HiFi (CCS) reads were generated with ccs v8.0.1, segmented into Kinnex S-reads with skera v1.2.0, demultiplexed/trimmed with lima v2.9.0, and refined to full-length non-chimeric (FLNC) reads using isoseq refine v4.2.0, after which reads were aligned to GRCh38 with pbmm2 v1.13.1. All steps up to and including the generation of aligned BAM files were run within SMRT Link v13.0.0.207600 (PacBio’s web-based, end-to-end workflow manager) and were executed per sample. After SMRT Link completed per-sample alignment, the aligned BAMs from all samples were merged with samtools v1.20 to produce a single multi-sample BAM ^59^. Isoseq collapse was then run on this merged BAM with default settings. The resulting read_stat.txt contains the mapping between collapsed transcript IDs (e.g., PB.1.1) and unique movie names that encode the originating sample; extracting the movie name, mapping it to sample names, and tallying FLNC reads per PB ID yielded a PB-by-sample FLNC count matrix (Fig. S1).

Next, the collapsed isoforms were annotated with pigeon classify v1.3.0 (PacBio transcript toolkit; SQANTI3-based) using GENCODE V47 as the reference genome annotation, producing classification.txt, which contains metadata that were processed with pigeon filter v1.3.0 to remove isoforms flagged for likely internal priming and reverse-transcription switching artifacts, following default filters: --polya-percent 0.6, --polya-run-length 6, --max-distance 50, --mono-exon, --skip-junctions. ISM isoforms are often interpreted as partial fragments resulting from incomplete retrotranscription or mRNA decay, therefore, we only kept 5’ fragment isoforms that have 3’ support and 3’ fragment that have 5’ support from external datasets. 5’ support was based on CAGE peaks found in the refTSS v3.3 annotation^60^, while 3’ support was determined based on polyA site data from PolyASite 2.0 atlas^61^. Finally, for all downstream analyses we retained collapsed isoforms supported by >5 FLNC reads in more than two samples.

#### Short-read RNAseq data processing

Raw reads from short-read RNAseq were aligned to the GRCh38.p13 (hg38) reference genome via STAR v2.4.2a using comprehensive gene annotations from Gencode V47^62^. BAM files were produced in both genomic and transcriptomic coordinates and sorted using samtools v1.20^59^. Splice junctions from SJ.out.tab files produced by STAR were mapped to splice junctions in the long-read RNAseq dataset, with default parameters for splice junction output filtering in STAR.

#### Open reading frame prediction and classification

To predict open reading frames (ORFs) from the set of high-confidence full-length isoforms, we used the ORFanage tool^22^. We ran ORFanage 1.1.0 in BEST mode, using GENCODE V47 CDS as a reference, and denoted a novel isoform as a coding isoform if it was assigned an ORF by ORFanage. We used the default setting of 50 for the minimal number of nucleotides in an ORF for it to be included. Each unique ORF represents a unique full-length protein isoform. To systematically characterize these predicted protein isoforms, we adapted SQANTI protein^23^, a protein isoform classification scheme that is based on SQANTI3 transcript isoform classification scheme. Predicted protein isoforms are classified as either pFSM, pISM, pNIC, or pNNC.

#### Proteomics data analysis

The proteomics data were analyzed using a two-pronged strategy to enable both the quantification of known proteins and the identification of novel protein isoforms.

##### Quantification of known proteins

During the TMT-MS run, MS2 spectra were searched in real-time by the acquisition instrument against the UniProt human reference proteome (UP000005640)^63^. This was performed using a Sequest-based platform with the COMET algorithm^64^. Peptide-spectrum matches (PSMs) that passed a 1% false discovery rate (FDR) filter were selected for MS3 analysis. The quantification of known proteins was derived from the TMT reporter ion intensities in these MS3 spectra. Only peptides with an isolation specificity ≥ 0.5 and a summed signal-to-noise (S/N) ratio greater than 110 across all TMT channels were included in the final quantitative analysis. This pipeline provided robust quantification for the established proteome.

##### Identification of novel peptides

To identify peptides derived from the novel mRNA isoforms discovered in our long-read RNAseq data, we performed a separate, offline analysis guided by recent long-read proteogenomics approaches^23^. All acquired MS2 spectra were searched using Comet version 2024.01 rev. 1 against a custom protein sequence database^64^. This database was constructed by combining all protein-coding sequences from the GENCODE V47 reference with the predicted amino acid sequences of our novel isoforms. The search parameters included methionine oxidation as a variable modification, and static modifications for cysteine carbamidomethylation and TMT labeling of peptide N-termini and lysine residues.

Following the search, peptide spectral matches (PSMs) were pooled and re-scored using Percolator (v3.05.0) to improve identification accuracy^65,66^. Only PSMs with a re-scored FDR-corrected *P*-value < 0.05 were retained. We mapped identified peptides to their potential parent isoforms using a custom Python script that enforces tryptic digestion rules and accounts for isoleucine/leucine isobaric equivalence. Following this, a peptide was classified as “novel” if it mapped uniquely to one of our novel protein isoform sequences and not to any entry in the GENCODE reference and “known” otherwise. Annotated spectra were visualized and exported for display using the Interactive Peptide Spectral Annotator^67^.

Since these novel peptides were identified solely through offline re-analysis of MS2 spectra and were not prioritized for MS3 fragmentation during the instrument’s real-time acquisition cycle, TMT-based quantification was not possible. Therefore, this pipeline serves to validate the existence of these novel proteins within the multiplexed pool, but cannot resolve their sample of origin or quantify their expression levels.

#### Ribosome profiling reanalysis

In brief, Ribo-seq involves treating mRNAs with ribonuclease to degrade the regions that are not protected by the ribosome, leaving behind ∼30 nt fragments, known as ribosome protected mRNA fragments. These fragments are mapped back to the original mRNA to define actively translated regions^68^.

Ribosome profiling data from Duffy et al. were sequentially trimmed to remove adapters and poly(A) sequences using *cutadapt*^25,69^. Reads were processed with the following commands:

1. *cutadapt -j 20 -a AGAGCACACGTCTGAACTCCAGTCACX -O 2 -m 14*
2. *cutadapt -a AGATCGGAX -O 1 -e 2 -m 14*
3. *cutadapt -a A{10} -O 3 -m 14*

Trimmed reads were aligned to the human genome using STAR^62^, allowing a maximum of two genomic alignments per read (*--outFilterMultimapNmax 2*). The STAR command used was:

*STAR --runThreadN 6 --runMode alignReads --outFilterMultimapNmax 2 --outFilterType BySJout --outSAMattributes All --outSAMtype BAM SortedByCoordinate --readFilesCommand zcat*

Mapped reads were quantified with *HTSeq-count*^70^. Regions for quantification included annotated coding and untranslated regions from GENCODE V47, as well as novel coding and untranslated regions predicted from our long-read sequencing data using ORFanage^71^. To count only uniquely mapping reads fully contained within each region, we used:

#### htseq-count -m intersection-strict -f bam

Following quantification, regions were required to have at least six RNAseq reads in the matched input libraries to be retained for downstream analysis. Translational efficiency (TE) was calculated by adding a pseudocount of 0.5 to all measurements, dividing ribosome-protected fragment counts by matched input RNAseq counts for each region, averaging TE across the three biological replicates, and log2-transforming the resulting values. Statistical comparisons of TE between predicted coding and untranslated regions were performed using the Wilcoxon rank-sum test (*wilcox.test*, default parameters in R).

#### Analysis of Differential Gene and Transcript Expression

##### Preprocessing and Differential Gene Expression (DGE)

For all differential transcript and gene analyses, we used transcripts from our long-read RNAseq data with predicted ORFs (according to ORFanage) in the FSM, ISM, NIC or NNC categories by SQANTI (n = 158,844 transcripts; 14,104 genes). All gene-level analyses were performed using official gene names rather than PacBio IDs (PB IDs), as PB IDs can ambiguously map to numerous genes across a genomic region. Gene expression was calculated as the sum of transcript expression for all isoforms attributed to a given gene. The raw transcript count data were processed and normalized using the edgeR package^72^ to generate counts per million (CPM) values via the cpm() function.

Differential gene expression (DGE) was calculated between timepoints (t00 vs. t04, t04 vs. t30, and t00 vs. t30) using DESeq2^73^. A positive log2 fold change in these comparisons reflects an increase in expression from the earlier to the later timepoint.

##### Differential Transcript Usage (DTU) and Expression (DTE)

To analyze isoform-specific dynamics, we imported isoform count data, CPM values, and exon structure into *IsoformSwitchAnalyzeR*^27,74^ using the *importRdata()* function with the detectUnwantedEffects argument set to FALSE. We then performed a stringent pre-filtering step using the prefilter() function with the following parameters: *geneExpressionCutoff = 1 CPM*, *isoformExpressionCutoff = 0*, and *IFcutoff = 0.01*. This filtering retained 68,801 isoforms where the parent gene was expressed and the isoform constituted at least 1% of the gene’s total expression in at least one condition.

Using this filtered set, we performed two parallel analyses. First, Differential Transcript Usage (DTU) was calculated using the *isoformSwitchTestDEXSeq()* function. Switches were considered significant if the absolute change in isoform fraction (|ΔIF|) was > 0.05 with an FDR < 0.05. Second, Differential Transcript Expression (DTE) was calculated on the exact same set of 68,801 transcripts using DESeq2. Transcripts were considered differentially expressed if they had a |log2FC| > 1 and FDR < 0.05.

##### Functional Analysis of Isoform Switching

To characterize the biological impact of isoform switching, we performed a deep functional annotation. Alternative splicing events were identified using *spliceR*^75^ by comparing each isoform to a hypothetical pre-mRNA. Microexons were specifically defined as skipped exons less than 28 nucleotides in length, following prior definitions^3^. We further annotated transcripts for features such as retained introns (12,224 transcripts) and premature termination codons (PTCs; 7,343 transcripts), with a PTC defined as a stop codon located ≥50 nucleotides upstream of the final exon-exon junction.

To predict consequences at the protein level, we integrated annotations of protein domains using Pfam^76^, and disordered regions using IUPred2A^77^. We then used the *extractConsequenceEnrichment()* and *extractSplicingEnrichment()* functions in IsoformSwitchAnalyzeR to test for the enrichment of specific splicing events and functional consequences among switching isoforms (alpha < 0.05, dIF > 0.05), summarizing results at the gene level. For this analysis, a minor custom modification was made to the compareAnnotationOfTwoIsoforms() function to ensure correct identification of transcription termination sites (TTSs). Finally, microexons identified in significant isoform switches were cross-referenced with the vastDB database^21^ and GENCODE V47 by their genomic coordinates.

###### Analysis of Differential Protein Expression

Differential protein expression was calculated using MSstatsTMT^78^. The intensity of identical peptides in the same channel were summed. Protein expression was normalized across TMT channels using *proteinSummarization*. Differential protein expression for each timepoint comparison was calculated using *groupComparisonTMT*.

###### Integrated Clustering of Expression Dynamics

To categorize and compare the dynamic trajectories of genes and proteins, we classified their expression patterns into one of nine clusters. The classification used a two-character code representing the expression change from t00-t04 (first character) and t04-t30 (second character). The symbols ‘U’ (increase; log2FC > 1; FDR < 0.05), ‘D’ (decrease; log2FC < -1; FDR < 0.05), and ‘–’ (no change; |log2FC| <1) were assigned based on each respective threshold. This method was applied independently to both the gene-level RNA and protein expression data.

#### Gene Set and Functional Enrichment Analysis

Gene Ontology (GO) term^79^ enrichment was conducted using the *gprofiler2* package^80^. Enriched “GO:BP”, “GO:MF”, and “GO:CC” terms were determined for the nine protein expression trajectories, with the background set to all proteins identified in our mass spectrometry analysis. Benjamini-Hochberg corrections were used for multiple comparisons and GO terms with adjusted *P*-values <0.05 were deemed significant. ASD risk gene enrichment was tested using all genes present in SFARI Gene (Q1 2025). For each protein expression cluster, ASD risk gene enrichment was tested using a one-sided Fisher’s exact test, with all expressed proteins as the background.

For both ASD risk gene and GO term enrichment analyses of isoform switch consequences, genes with at least one isoform switching event resulting in either direction of the event (e.g., exon skipping or exon inclusion) were used. The background genes were all genes in the *IsoformSwitchAnalyzeR* analysis (n = 10,529). GO term^79^ enrichment was conducted using the *gprofiler2* package^80^. Enriched “GO:BP”, “GO:MF”, and “GO:CC” terms were determined. Benjamini-Hochberg corrections were used for multiple comparisons and GO terms with adjusted *P*-values <0.05 were deemed significant. For each event, a one-sided Fisher’s exact test to determine if there was significant enrichment between the genes with each event, and all SFARI genes, with the background genes as all genes in the *IsoformSwitchAnalyzeR* analysis (n = 10,529). Bonferroni test correction was applied within the statistical tests performed for each timepoint comparison.

#### Identification and Characterization of AS-APA Coordination

To investigate the coordination between alternative splicing (AS) and alternative polyadenylation (APA), we developed a novel statistical approach using generalized linear models using quasi-binomial models. We felt that this was necessary because established methods, such as those using contingency tables, typically require pooling data across replicates, which can obscure sample-to-sample variance. Our approach was designed to overcome these limitations by explicitly modeling this sample-to-sample variance and time as variables and we performed a rigorous comparison between our novel and a more traditional approach.

##### Quasi-binomial approach

To investigate the coordination between exon splicing and APA, we performed a custom analysis on all 158,844 predicted protein-coding transcripts belonging to the FSM, ISM, NIC, or NNC categories. First, we identified AS (alternatively spliced) exons using the generateEvents function in SUPPA2^81^. Concurrently, we defined distinct polyA sites by clustering transcript end sites located within 24 nucleotides of each other. The initial dataset was filtered for isoforms containing at least one AS exon and genes with more than one distinct polyA site.

To prepare the data for statistical analysis, we quantified sample-specific read counts for exon inclusion and skipping at each distinct polyA site. We applied stringent filtering to retain robust events: each polyA site was required to have a cumulative (inclusion or skipping) count of ≥50 reads in at least five samples, and a cumulative count of ≥10 reads in each of those samples. Events with at least two polyA sites passing this filter were retained. A pseudocount of 1 was added to all counts to prevent zero-count errors in downstream calculations.

For each splicing event, we fit statistical models using the glm() function in R (with family = quasibinomial) to estimate the association of time point and polyA site usage on exon inclusion levels. We chose to use the quasibinomial family here to accommodate potential overdispersion in splicing read counts. We fit a series of three nested models per AS event:

1. Inclusion ∼ Time (Reduced model)
2. Inclusion ∼ Time + PolyA_Site (Global model)
3. Inclusion ∼ Time x PolyA_Site (Interaction model)

Both Time (t00, t04, t30) and PolyA_Site were treated as discrete variables. Events with non-convergent models were removed.

We performed model comparisons using F-tests on analysis of deviance (ANOVA) tables to compute likelihood ratio test *P*-values, which were then corrected using the Benjamini-Hochberg (FDR) method. Global effects (a stable association between polyA site and exon inclusion/exclusion) were evaluated by comparing Model 2 to Model 1. Interaction effects (a timepoint-dependent association) were evaluated by comparing Model 3 to Model 2. For each event, estimated marginal means were computed using the emmeans() function from the emmeans R package^82^ for Time within each PolyA_Site. The resulting emmean values represent the predicted probability of inclusion, along with 95% confidence intervals.

To ensure biological significance, we applied stringent effect size filters to all FDR-significant events, using the observed PSI values for each polyA site and timepoint. For the global events to be deemed significant, we required that the ΔPSI (maximum PSI - minimum PSI) between the polyA sites, averaged across time, had to be ≥0.1. An interaction event required two conditions to be met, demonstrating that the coordination was both time-dependent and had a substantial effect. First, the change in PSI across time for at least one polyA site had to be ≥0.1. Second, the difference in PSI between polyA sites at a single timepoint had to be ≥0.1. Global and Interaction coordination events were only considered significant if they passed both the statistical (FDR) and these effect size (ΔPSI) thresholds.

For each significant event, we assessed three metrics. First, we calculated the genomic distance between the 5’ position of the exon immediately downstream of the coordinated AS exon to the most proximal polyA site. Second, we counted the number of exons between the coordinated exon (exclusive) and the most proximal polyA site (inclusive), using the most abundant transcript as a representative transcript for the gene. Lastly, we annotated the genomic context of each coordinated exon (5’UTR, ORF, or 3’UTR) based on its location within its most abundant isoform. “5’ UTR + ORF” refers to exons with both 5’UTR and ORF annotations. “ORF + 3’UTR” refers to exons with both ORF and 3’UTR annotations.

We used a one-sided Fisher’s exact test to determine if there was significant enrichment between the genes with a significant coordination event, and all SFARI genes, with the background set of genes defined as all genes that were tested in this analysis.

##### Fisher’s exact test approach

To validate our quasi-binomial method for assessing coordination between exon AS and APA, we applied an alternative method used by previous reports^10,32^, but that does explicitly consider sample-level variance and relies on pooling data across replicates. We used the same exon splicing and polyA determination described above.

We then constructed a primary contingency table for each exon splicing event, creating an n x 2 matrix where the n rows represented the polyA sites and the two columns contained the counts for exon inclusion and exclusion. To ensure statistical power and remove noisy, low-count events, we applied a series of stringent quality control filters. First, we removed any polyA site with <100 total counts (across exon inclusion and exclusion). Second, we removed any splicing event where the total counts for either the included or the skipped isoform were <100. After filtering, events with >1 polyA site were kept. A pseudocount of 1 was then added to all cells in the remaining matrices to avoid issues with zero counts in downstream calculations.

With the filtered data, we then proceeded to the formal statistical test for coordination. To assess the influence of each individual polyA site on the splicing outcome, we iteratively tested each site against all others. For each polyA site within a given splicing event, we generated a 2×2 contingency table comparing the inclusion/exclusion counts for that specific site against the pooled counts from all other polyA sites for the same event. A Fisher’s exact test was applied to each of these 2×2 tables to determine if the choice of that particular polyA site was significantly associated with the splicing outcome. The resulting *P*-values were corrected for multiple comparisons using the Benjamini-Hochberg procedure. To quantify the effect size, odds ratios (OR) were log-transformed, with a minor constant (1 x 10⁻¹³) added to prevent computational errors. A significant coordination event was ultimately defined as an event containing at least one polyA site that satisfied two criteria: an FDR < 0.05 and an absolute log odds ratio (|logOR|) of 1 or greater.

To compare these results with those from the quasi-binomial modelling approach, we determined the set of events tested in both analyses (n = 3,169 events). We compared significant Fisher’s exact test events with significant global events. Out of 2,636 total significant events, 1,704 (65%) were overlapping.

###### Analysis of Alternative Polyadenylation (APA)

To quantify dynamic changes in polyadenylation (polyA) site usage, we first processed all 158,844 protein-coding transcripts using the qapa build function from QAPA^34^, skipping the annotation step. After filtering for genes with more than one polyA site and expression >1 CPM at each timepoint, we calculated the relative usage of each 3’UTR (PolyA site Usage, PAU) as the ratio of the UTR’s CPM to the total CPM for the parent gene. For each timepoint, we calculated median PAU. For each gene, we designated the PAU value from the most proximal polyA site as PPAU. The change between timepoints (ΔPPAU) was calculated as PPAU(early) - PPAU(late), with a ΔPPAU > 20 defined as a 3’UTR lengthening event and a ΔPPAU < -20 as a shortening event.

To assess the relationship between these APA dynamics and gene expression, we calculated the Pearson correlation coefficient between each gene’s ΔPPAU and its log2 fold change as determined by DESeq2^73^, only using transcripts with a UTR in the QAPA analysis.

To isolate the effect of APA from other upstream changes in transcript structure, we performed a secondary analysis. We first clustered transcripts into groups that shared an identical sequence up to the beginning of the 3’UTR. Within these clusters, distinct polyA sites were defined by grouping transcript end sites located within 24 nucleotides. These sequence clusters were then treated as independent ‘genes’, and the quantification and correlation analysis described above was repeated at the cluster level.

### Mutational Constraint and ASD Variant Burden Analysis

#### Data Resources and Variant Filtering

The mutational constraint analysis was performed on WES data from 730,947 individuals and WGS data from 76,215 individuals in gnomAD v4.1, the generation and quality control of which have been previously described^37,38^. We retained only variants passing all three filtering criteria computed by the gnomAD team (VQSR, AC0 and InbreedingCoeff).

The ASD burden analysis was performed using the MSSNG, SSC, and SPARK cohorts, comprising WES or WGS of ASD individuals, unaffected siblings, and both parents^39–41^. Total cohort counts of ASD individuals and unaffected siblings with at least one parent sequenced are as follows: MSSNG (affected n = 6,006; unaffected siblings n = 526), SSC (affected n = 2,419; unaffected siblings n = 1,967), SPARK WES (affected n = 53,516; unaffected siblings n = 24,449) and SPARK WGS (affected n = 3,540; unaffected siblings n = 2,226). *De novo* single nucleotide variants (SNVs) and insertion/deletions (Indels) were extracted from individuals with ASD and their unaffected siblings who had both parents sequenced.

#### Variant Effect Prediction

Variant effects were predicted for QC-passing variants in gnomAD and *de novo* mutations in ASD cohorts using (1) the GENCODE V47 reference transcriptome, and (2) our custom reference transcriptome derived from long-read RNAseq, with protein-coding regions predicted by ORFanage. For the MAPS analysis, we used the Ensembl Variant Effect Predictor (VEP) v114^83^ to predict the effect of gnomAD variants. Known synonymous, missense and nonsense (i.e., stop-gain) variants are defined as variants that are predicted to be synonymous, missense and nonsense using the GENCODE reference transcriptome. Novel synonymous, missense and nonsense variants are defined as variants that are present in non-coding regions as defined by GENCODE V47, but predicted to be synonymous, missense and nonsense using our custom reference transcriptome. For the ASD burden analysis, we used ANNOVAR (2019Oct24 release)^84^ to predict the effects of *de novo* mutations in ASD cohorts, but predicted to be in a less damaging category using the GENCODE reference transcriptome. This strategy enabled the identification of mutations with potentially damaging effects that would be missed by standard annotations.

#### Mutational Constraint Analysis

To assess negative selection acting on known and novel transcriptomic elements, we calculated the Mutability-Adjusted Proportion of Singletons (MAPS) metric for known and novel variant classes^43,85^. MAPS identifies classes of variants that are depleted in the general population and are therefore likely to be deleterious. We trained a model using synonymous variants as the neutral baseline, adjusting for local sequence context by regressing the proportion of singletons against trinucleotide-specific mutation rates^86^. For near-splice regions, which were defined as ±10 nucleotides flanking a splice junction (i.e., the exon-intron boundary), variants were grouped by their position relative to the known or novel splice site. For variants in protein-coding regions, variants were grouped by their predicted consequence (e.g., missense, nonsense) based on either GENCODE or our custom ORFanage-based reference transcriptome.

#### ASD De Novo Variant Burden Analysis

For the *de novo* variant burden analysis, known splice site variants are defined as variants that intersect the intronic dinucleotides immediately upstream or downstream of exons in GENCODE reference transcriptome, whereas novel splice site variants are defined as variants that intersect intronic dinucleotides upstream or downstream of novel splice sites in our custom reference transcriptome only. To test for an enrichment of these variant classes in ASD, we used Fisher’s Exact Test to compare the proportion of individuals with at least one qualifying variant in the ASD group versus the unaffected sibling control group. The analysis was conducted for each cohort individually and as a combined analysis across all cohorts.

## Supplemental Figures

**Figure S1.**
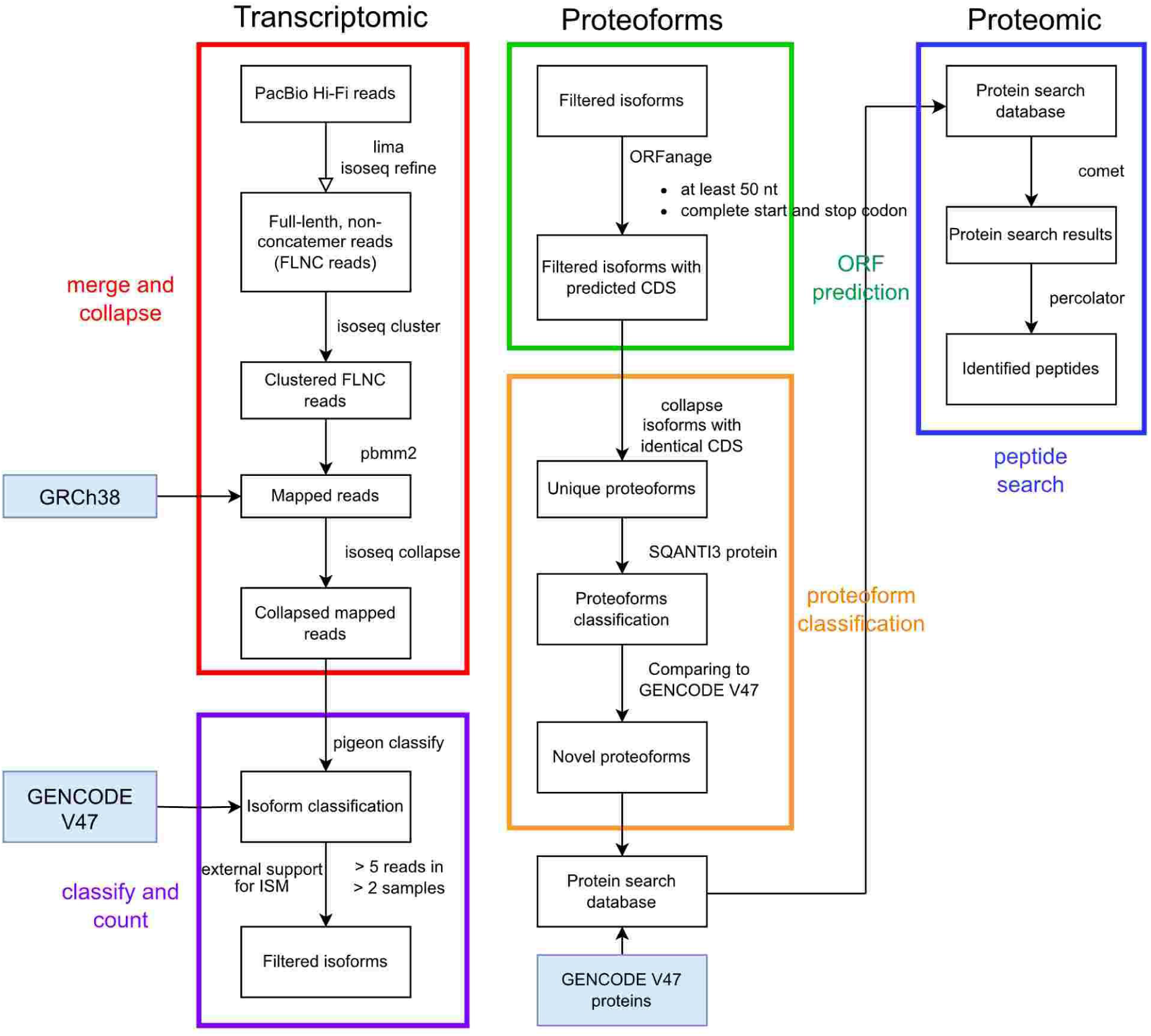
An overview of the proteogenomic bioinformatics pipeline. This flowchart details the processing of raw long-read transcriptomic data (red box) and its classification against GENCODE V47 to define a high-confidence set of filtered isoforms (purple box). This transcriptome is then used to predict open reading frames (ORFs) (green box) and classify novel proteoforms (orange box). These novel proteoforms are combined with reference proteins to create a custom database used for the proteomic peptide search (blue box) to validate translation. Further details are provided in Methods.

**Figure S2.**
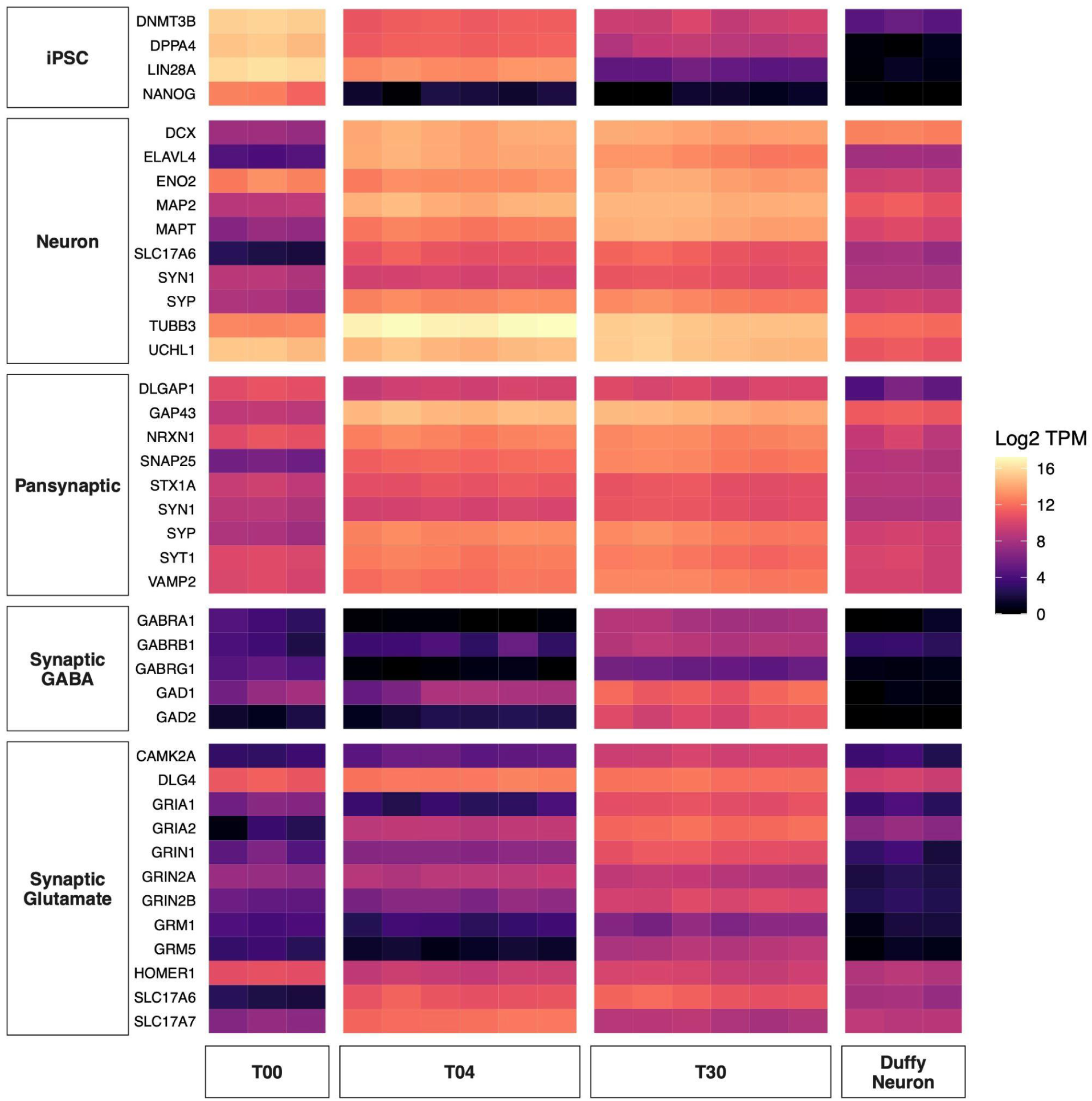
Expression of canonical cell-type markers confirms cellular identity across the differentiation time course. The heatmap shows gene expression (Log2 TPM) from this study’s short-read RNAseq data at t00 (iPSC), t04 (intermediate), and t30 (neuron) time points. The final column displays expression from an external NGN2-based iPSC-derived neuron dataset for comparison^25^.

**Figure S3.**
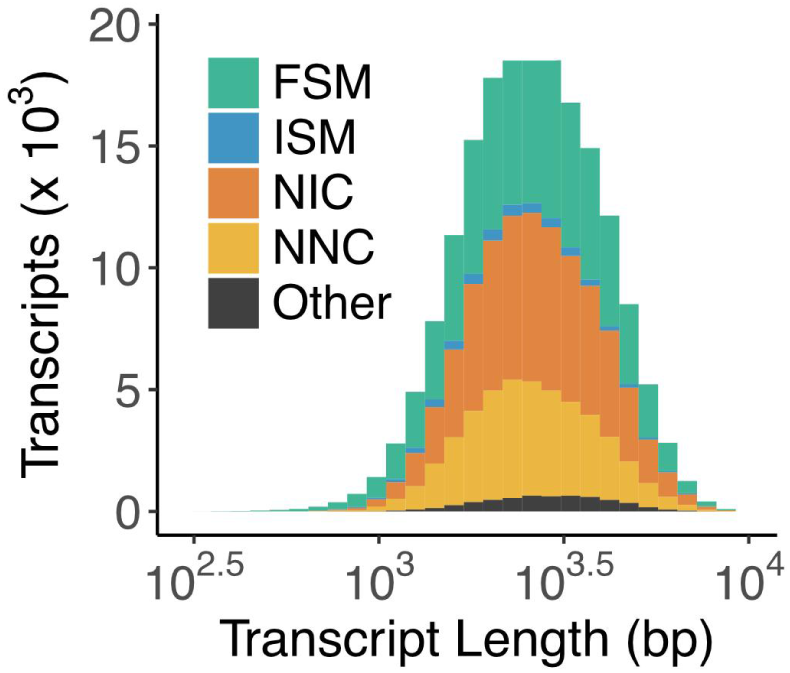
Distribution of isoform lengths from long-read RNAseq. Related to Fig. 1. This plot shows that isoforms of different structural categories have a similar distribution of transcript length that centres around 2600 bp.

**Figure S4.**
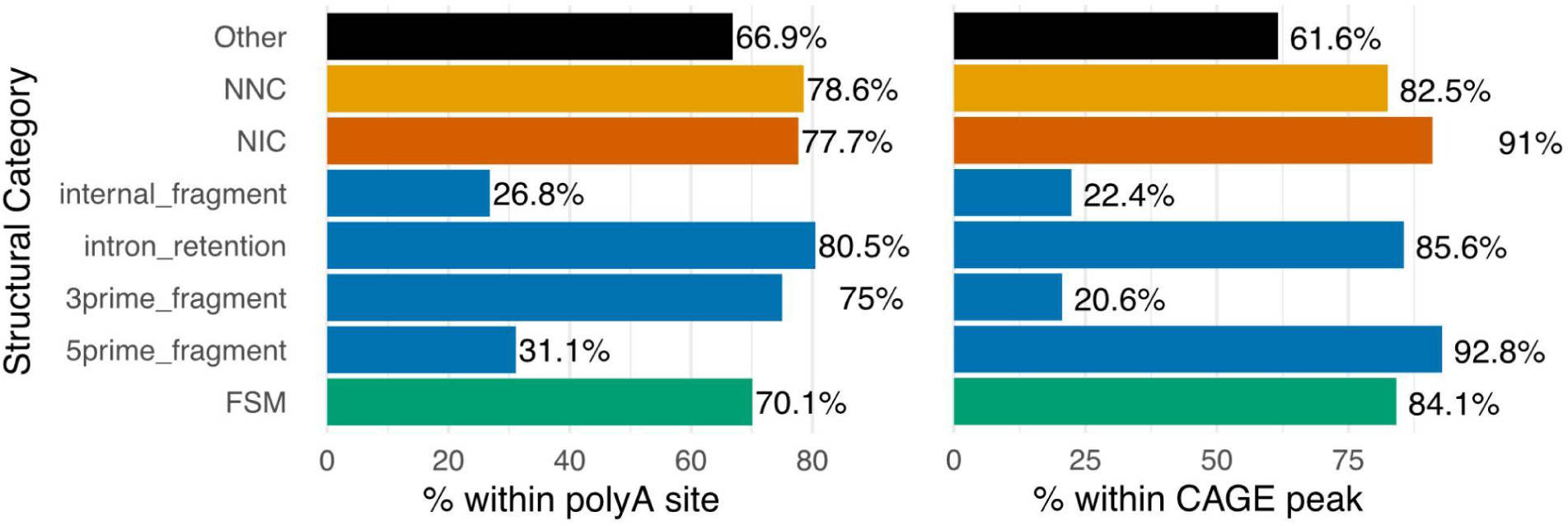
Validation of isoform 3’ and 5’ ends using external datasets. Related to Fig. 1. Bar plots show the percentage of isoforms whose 3’ ends overlap a known polyA site (left) and whose 5’ ends overlap a CAGE peak (right). Structural categories match Fig. 1, with Incomplete Splice Matches (ISM) further stratified into four subcategories: internal fragments, intron retention, 3’ fragments (putative 5’ truncation), and 5’ fragments (putative 3’ truncation). To minimize technical artifacts, only 3’ fragments with valid 5’ support (CAGE peaks) and 5’ fragments with valid 3’ support (polyA sites) were retained for the final high-confidence catalog.

**Figure S5.**
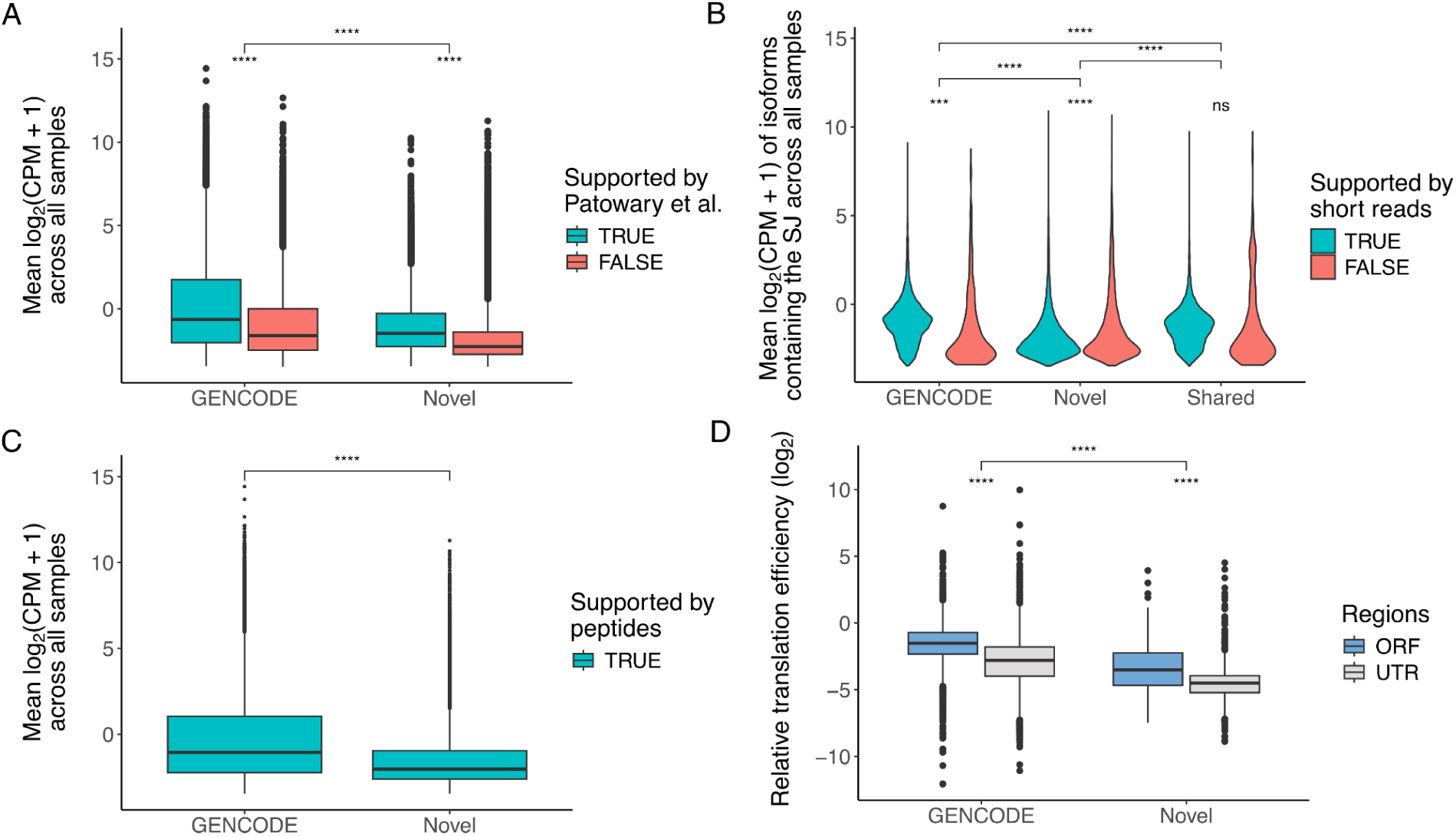
Transcript abundance is a major contributor to multi-omic validation of novel isoforms. **(A)** Mean expression (log2 CPM+1) of GENCODE and Novel isoforms, stratified by their detection in an external fetal brain long-read RNAseq dataset from Patowary et al., 2024. Isoforms replicated in Patowary et al have significantly higher expression than those unique to this study. **(B)** Mean expression of isoforms containing GENCODE, Novel, or Shared splice junctions, stratified by support from our matched short-read RNAseq. Validated junctions (teal) are associated with significantly higher isoform expression. **(C)** Mean expression of full-length isoforms supported by peptides from proteomics; where a peptide maps to multiple isoforms, expression is averaged across all isoforms containing peptide. Validated novel isoforms exhibit significantly lower expression than validated known isoforms. **(D)** Relative translational efficiency (TE), defined as the ratio of ribosome-protected fragments to input mRNA, calculated using external Ribo-seq data from human iPSC-derived neurons from Duffy et al., 2022. As a positive control, TE is significantly higher in known GENCODE coding sequences compared to 3’UTR. This pattern extends to the novel isoforms identified here, where predicted ORFs exhibit significantly greater TE compared to predicted 3’UTRs, providing independent validation for the ribosomal engagement and translation of these novel coding sequences. \*\*\**P* < 0.001; \*\*\*\**P* < 0.0001; ns = not significant.

**Figure S6.**
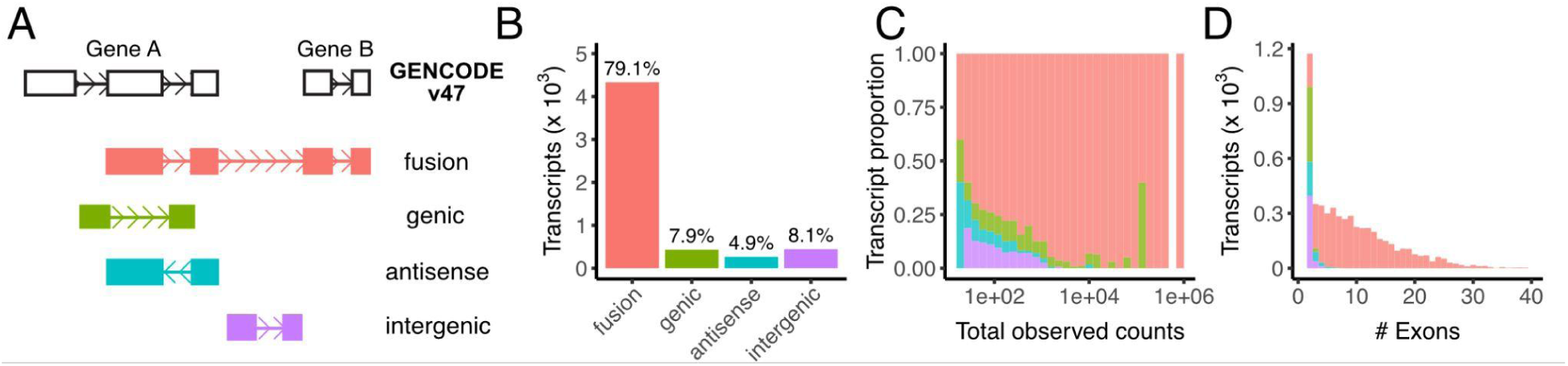
Classification and characterization of fusion, genic, antisense, and intergenic transcripts. (A) Schematic illustrating the classification of transcripts relative to the GENCODE V47 reference. Categories are defined as ’fusion’ (spanning two distinct gene loci), ’genic’ (overlapping a known gene), ’antisense’ (on the opposite strand of a known gene), and ’intergenic’ (in a previously unannotated region). (B-D) Characterization of these transcript categories. These structural categories are used to illustrate the total count and proportion of all high-confidence transcripts (B), as well as the distributions of total read counts (C) and exon counts (D) per transcript.

**Figure S7.**
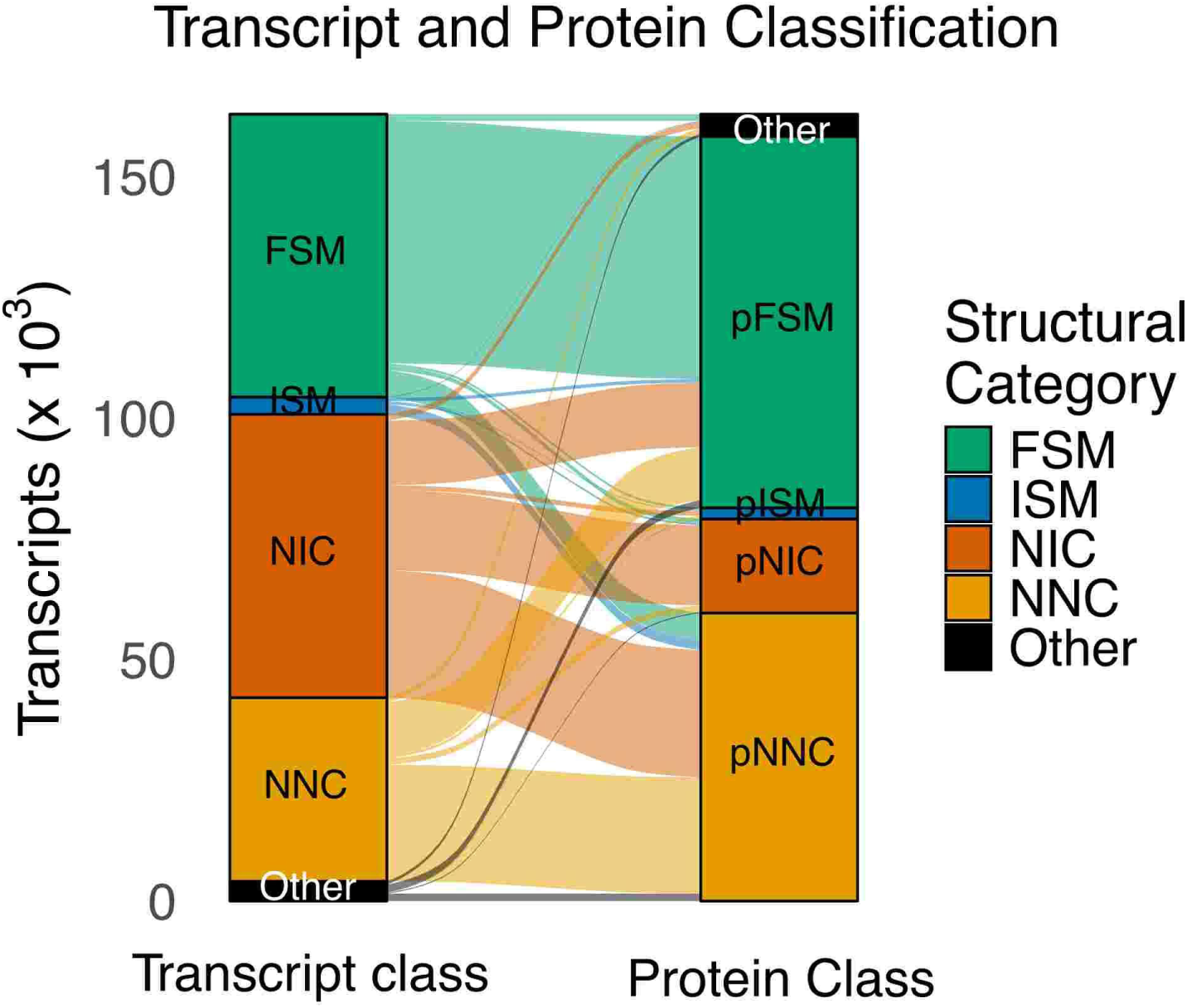
Relationship between transcript and protein structural classifications. Sankey diagram illustrating the mapping between the structural classification of mRNA isoforms and their corresponding predicted protein isoforms, using the SQANTI protein framework.

**Figure S8.**
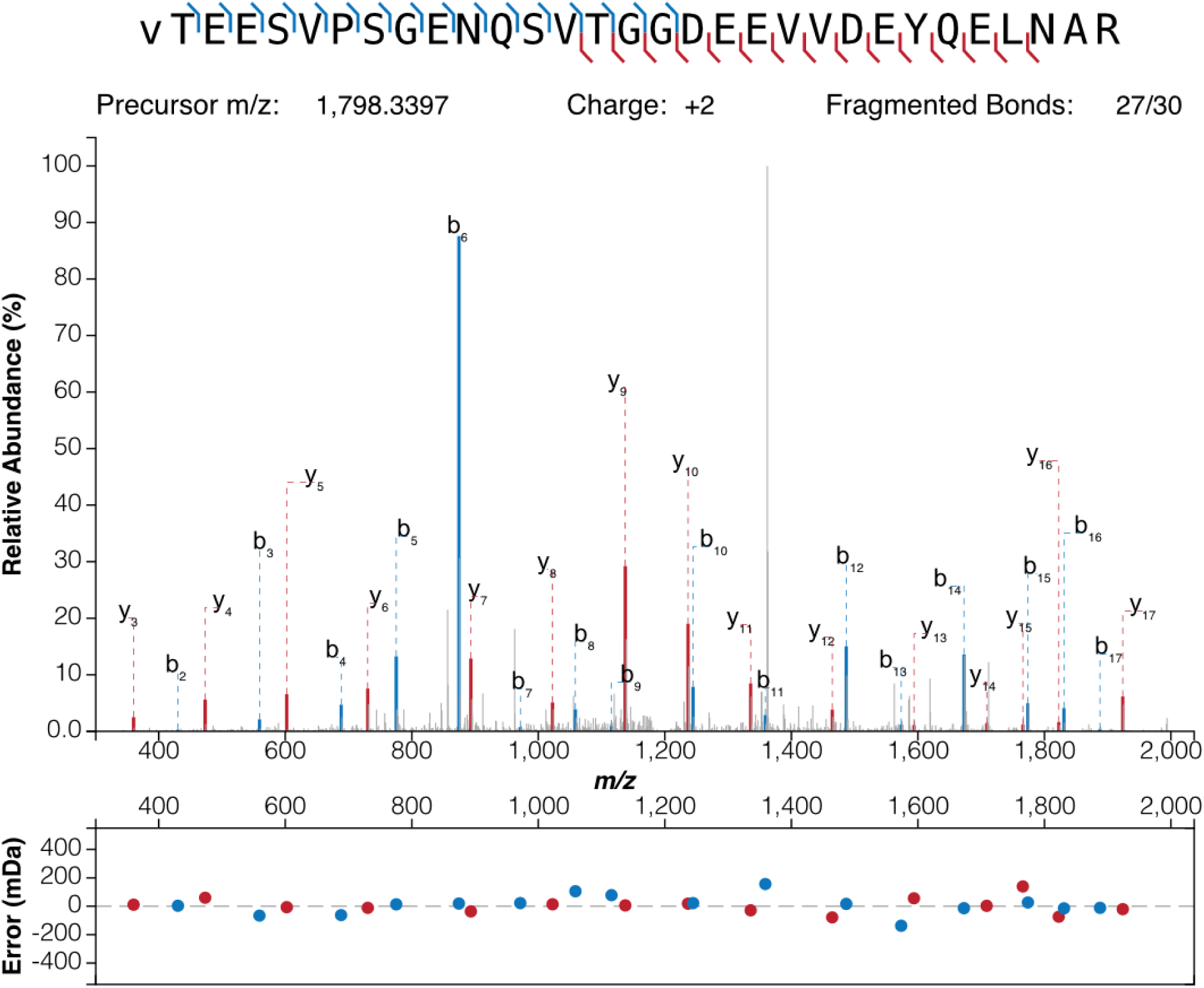
Peptide confirmation of the translation of a novel exon in EXOC1. Related to Fig. 2. Mass spectrum of peptide VTEESVPSGENQSVTGGDEEVVDEYQELNAR, which confirms the translation of the detected *EXOC1* microexon. Matched b and y ions are highlighted. m/z, mass/charge ratio.

**Figure S9.**
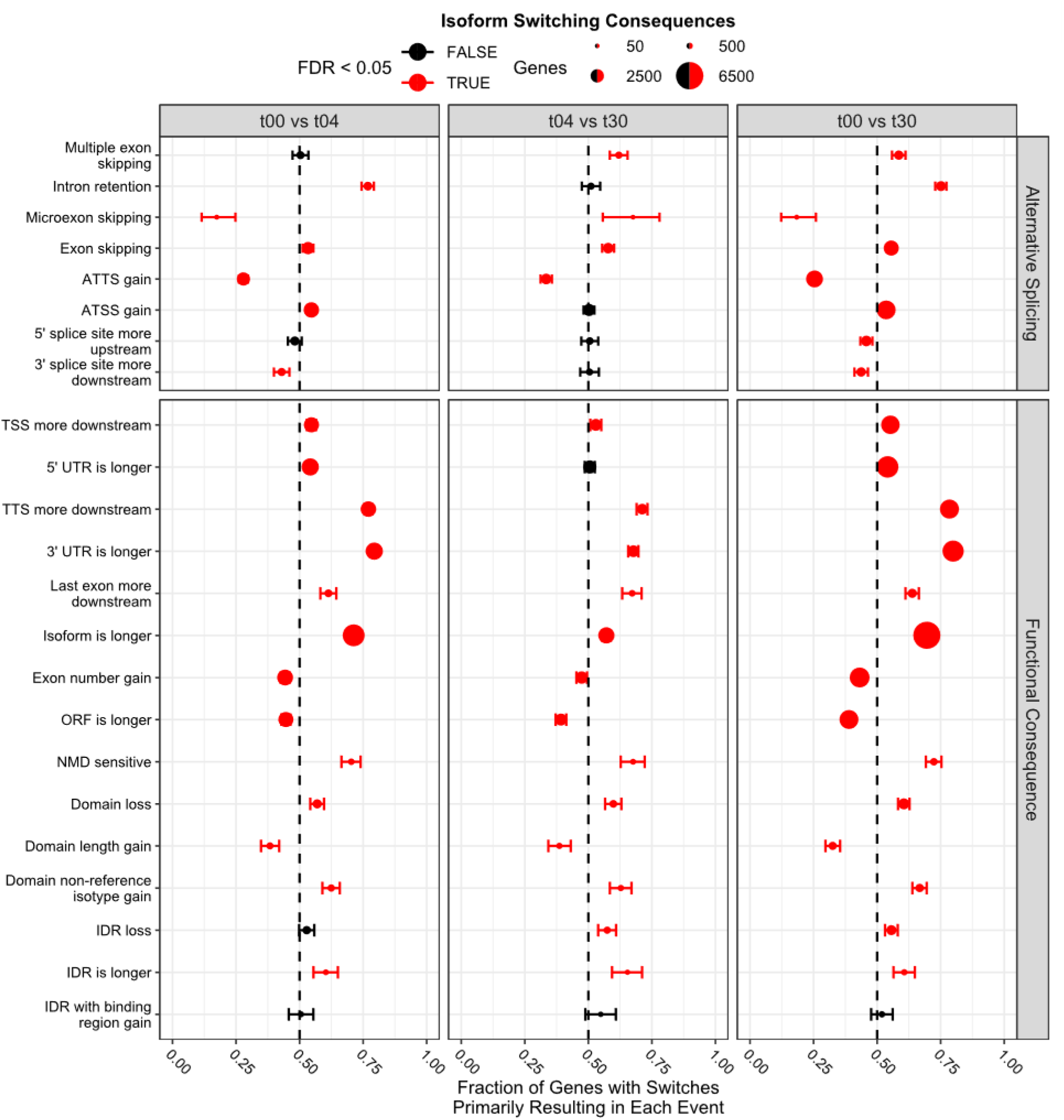
Complete list of isoform switching events for alternative splicing and functional consequences analyzed across differentiation. Points represent the fraction of genes where switches primarily result in a given outcome (e.g., exon skipping). The size of the points represents the total number of genes with isoform switches resulting in either opposing consequence. Error bars represent 95% confidence intervals; significance determined by binomial test.

**Figure S10.**
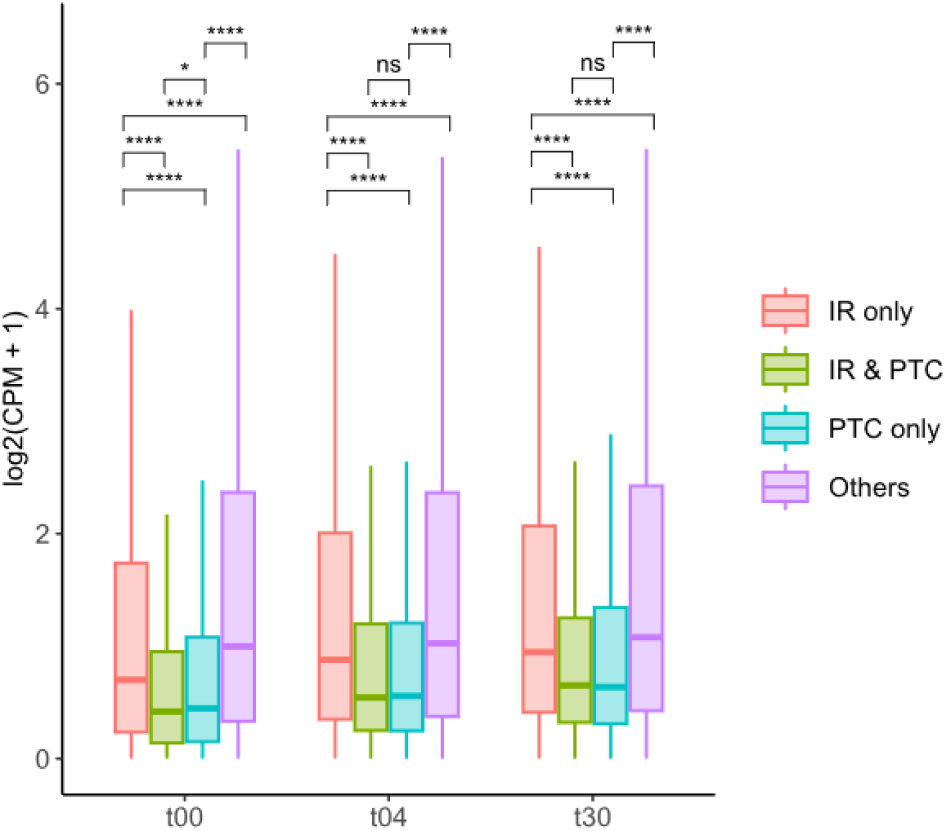
Expression of transcripts grouped by intron retention and premature termination codons. Boxplots compare the expression of transcripts as having Intron Retention (IR) only, IR and a Premature Termination Codon (PTC), a PTC only, and all other transcripts. Statistical comparisons within each time point were performed using a Wilcoxon rank-sum test with Bonferroni correction (\**P* < 0.05; \*\*\*\**P* < 0.0001; ns = not significant).

**Figure S11.**
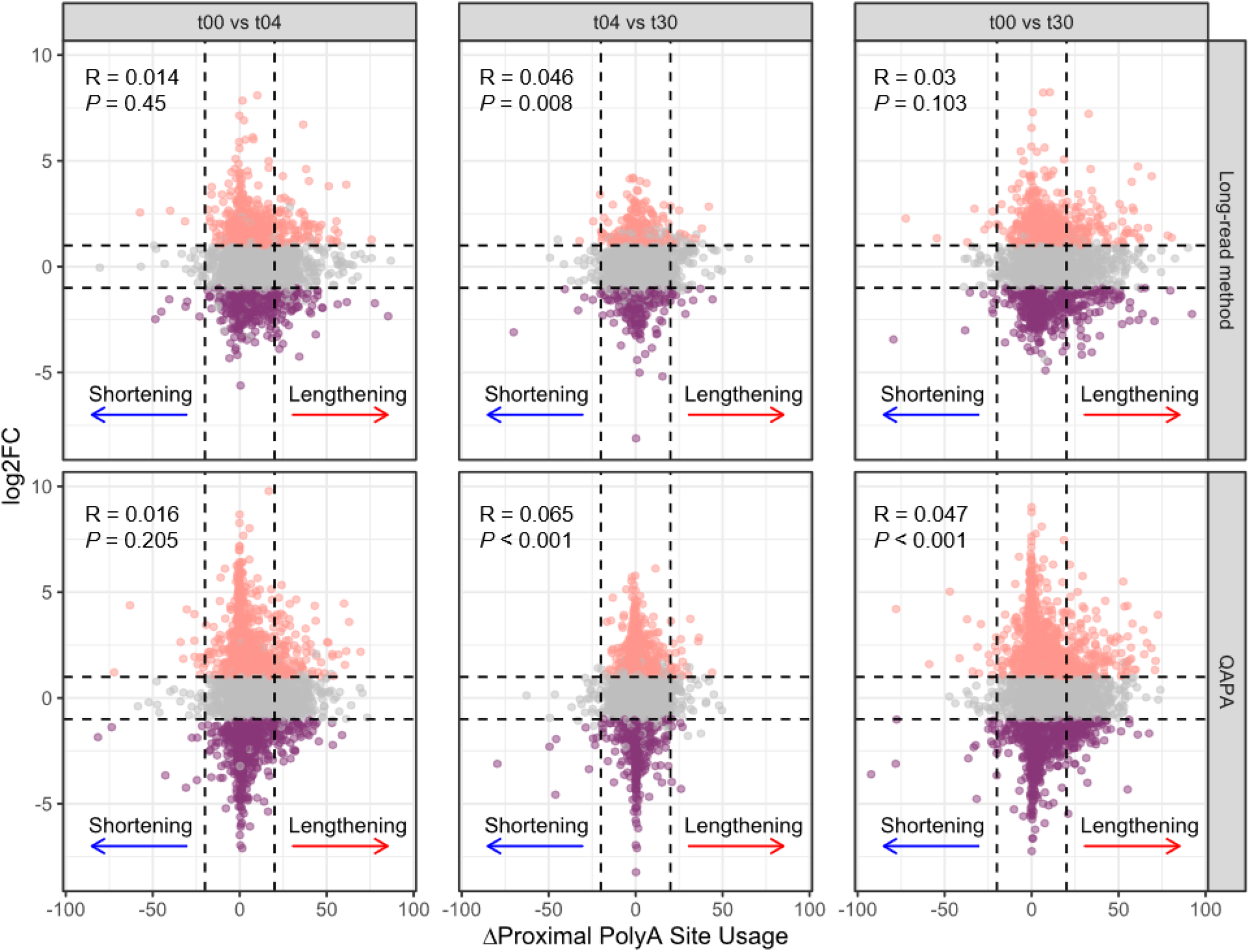
Lack of correlation between 3’ UTR length dynamics and mRNA expression changes. Scatter plots compare changes in gene expression (log_2_FC, y-axis) against the change in proximal polyA site usage (ΔPPAU, x-axis) between pairs of differentiation timepoints. Top row uses our custom long-read analysis method and bottom row uses QAPA for validation. 3′ UTR shortening (ΔPPAU < − 20, blue arrows) and lengthening (ΔPPAU > 20, red arrows) show no clear association with direction of gene expression changes (|log_2_FC| > 1, FDR < 0.05), including gene upregulation (peach dots) or downregulation (purple dots). Pearson correlation coefficients (R) and *P*-values for all datapoints are shown in each panel. Vertical dotted lines indicate ΔPPAU thresholds, and horizontal dotted lines indicate log_2_FC thresholds.

**Figure S12.**
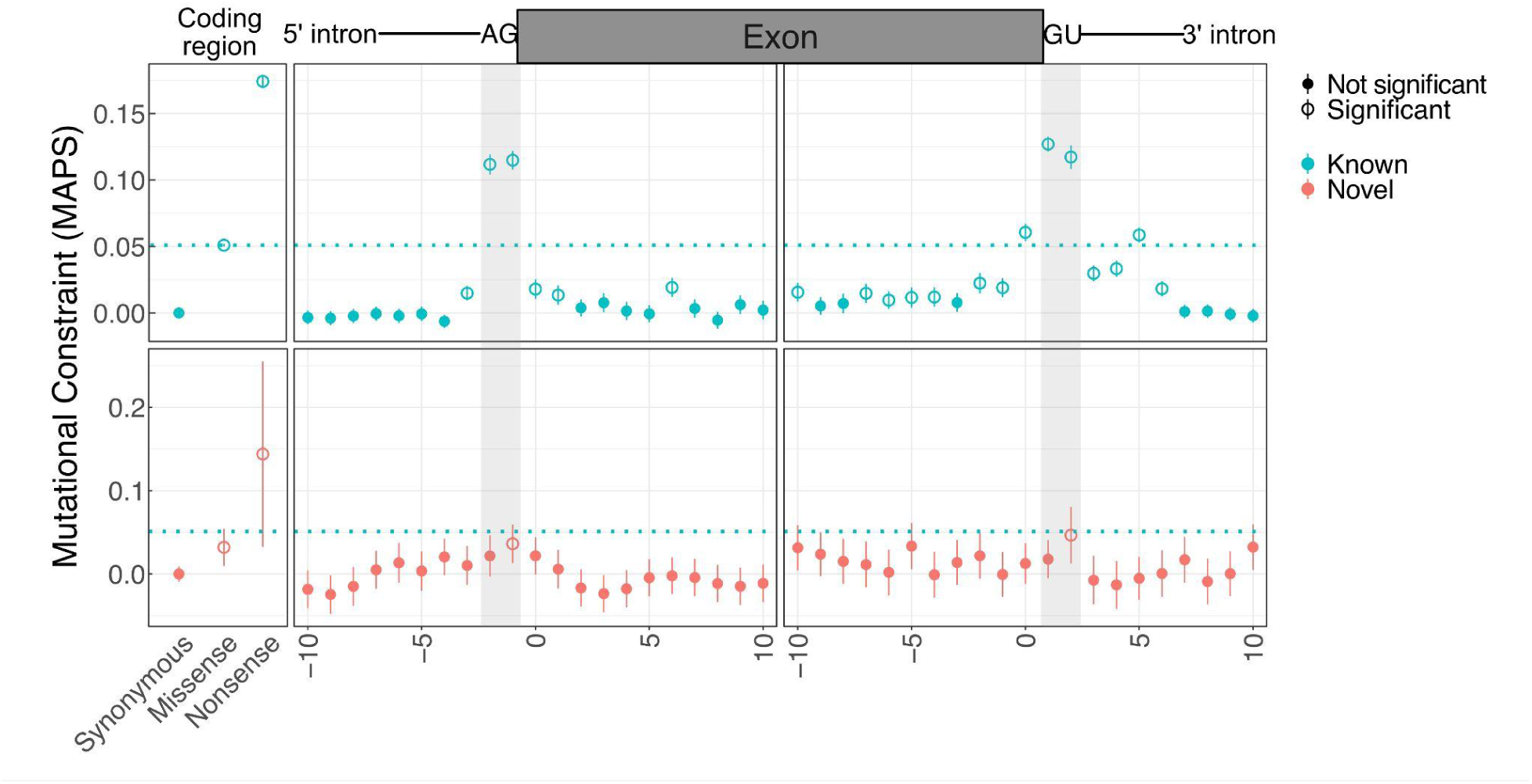
Mutational constraint of coding and near-splice variants using whole-genome sequencing (WGS) data. Related to Figure 6. The plots show mutational constraint (MAPS scores) for single nucleotide variants from 76,215 individuals in the gnomAD v4.1 WGS cohort. The top panel shows constraint at Known (GENCODE V47) elements, and the bottom panel shows constraint at Novel elements identified in this study. For coding consequences (left), scores are shown for synonymous, missense, and nonsense variants. Positions surrounding splice acceptor and donor regions are also shown, with shaded areas highlighting the two essential splice sites. Open circles indicate variant classes or positions with mutational constraint significantly different from synonymous variants (FDR < 0.05, chi-squared test). Dashed horizontal lines indicate the MAPS scores for known missense and nonsense variants, provided for context. Error bars represent 95% CIs.

**Figure S13.**
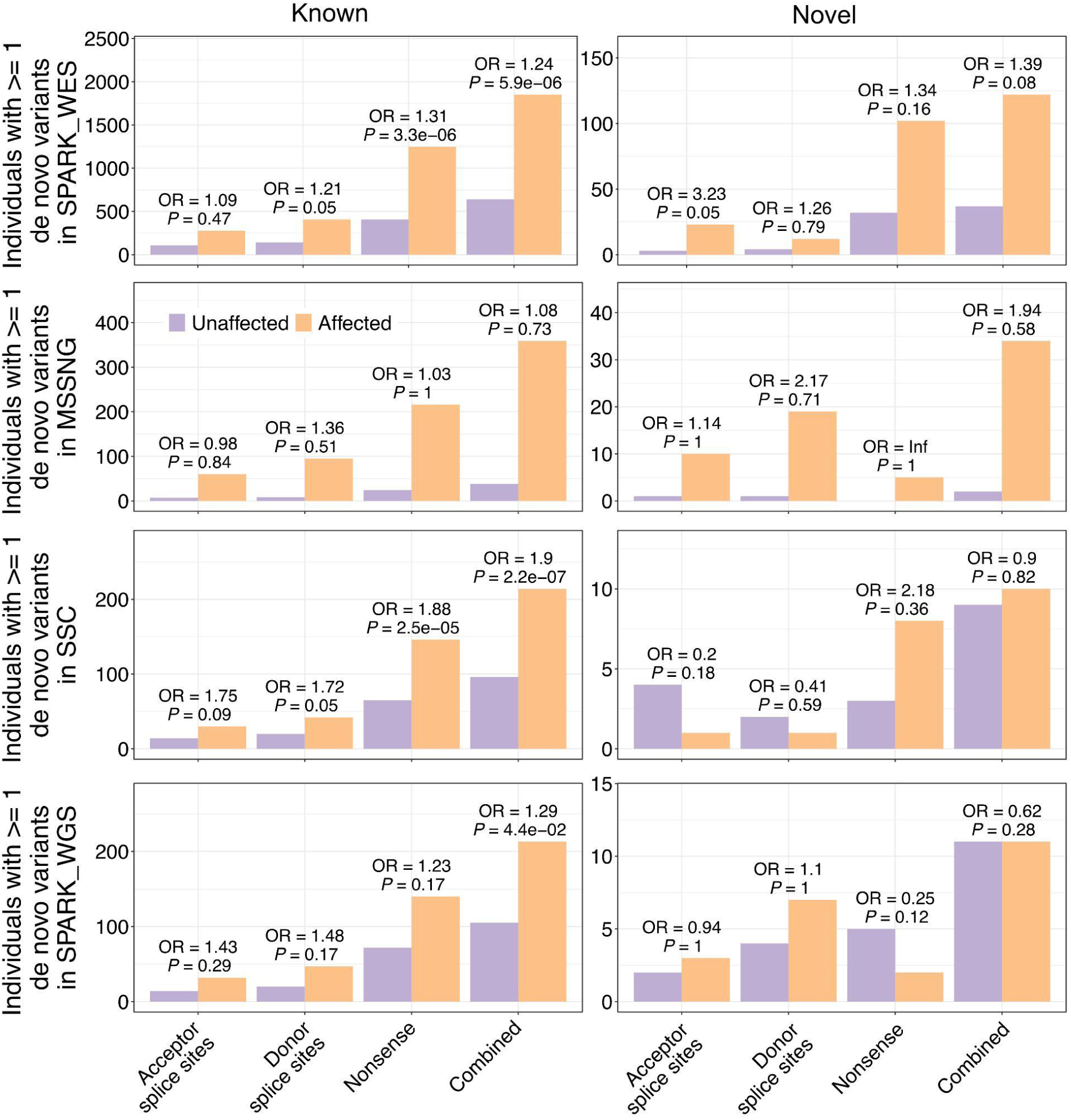
Burden of disruptive *de novo* mutations in individual ASD cohorts. Related to Figure 6. These plots show the burden of *de novo* mutations in individuals affected by versus unaffected by ASD for known (left) and novel (right) genomic elements. Each row represents data from one ASD cohort. Odds ratios (OR) and *P*-values from a Fisher’s exact test are shown.

## Supplementary Tables

**<u>Supplementary Table 1</u>**

**<u>Supplementary Table 2</u>**

**<u>Supplementary Table 3</u>**

**<u>Supplementary Table 4</u>**

**<u>Supplementary Table 5</u>**

**<u>Supplementary Table 6</u>**

**<u>Supplementary Table 7</u>**

**Table S1. Isoform Atlas and Annotation. Related to Figure 1**. This table is derived from classification.txt produced by pigeon classify, which is based on SQANTI3 QC. It is a tab-separated file where isoforms are rows and QC features computed by pigeon classify are columns. For a full glossary of columns and their meaning, see https://github.com/ConesaLab/SQANTI3/wiki/Understanding-the-output-of-SQANTI3-QC#classifcols. It contains 182,371 high-confidence mRNA isoforms identified across the t00, t04, and t30 time points. The table also includes an additional column that indicates the results of the overlap comparison with the external fetal brain dataset ^13^, where Ensembl or TALON IDs are listed if the isoform identified in our dataset overlaps with one in the Patowary dataset.

**Table S2. Catalog of Microexons identified in human iPSC-derived neurons.** This table lists the genomic coordinates and classification of 1,930 microexons in 8,008 transcripts (defined as exons < 28 nucleotides) detected in the long-read RNAseq dataset.

● **seqnames, start, end, strand**: Genomic location of the microexon (GRCh38).

● **width**: Length of the microexon in nucleotides.

● **gene_id/transcript_id**: Unique identifier for the PacBio gene ID and transcript isoform containing the microexon.

● **MIC_coord**: Combined genomic coordinates (chr:start-end).

● **GENCODE_v47**: Logical column indicating whether the microexon is found in GENCODE v47 (TRUE), or not (FALSE).

● **vastDB:** Logical column indicating whether the microexon is found in vastDB (TRUE), or not (FALSE).

● **EVENT:** vastDB exon ID.

**Table S3. Proteogenomic Validation of Novel Isoforms. Related to** Figure 2. This table in .csv file format, lists novel peptides identified by mass spectrometry that provide direct experimental evidence for the translation of protein-coding sequences. It provides the amino acid sequence for each peptide, the corresponding isoform ID(s), and the parent gene.

**Table S4. Gene and Protein Expression Dynamics. Related to** Figure 3. This table contains the complete gene-level differential expression results and functional enrichment analyses underlying the mRNA-protein dynamics shown in Figure 3.

● **Differential Gene Expression:** Full statistical results from DESeq2 for mRNA-level differential expression between time points (t00, t04, t30).

● **Differential Protein Expression:** Full statistical results from MSstatsTMT for protein-level differential expression.

● **Clusters:** The nine-category expression trajectory assignment (e.g., ’UU’, ’D-’) for each of the 8,498 genes with both mRNA and protein data.

● **SFARI Gene Enrichment:** Statistical results (odds ratio, *P*-value) for the enrichment of SFARI ASD risk genes in each protein and mRNA cluster, as shown in Figure 3D.

● **GO Term Enrichment:** Complete Gene Ontology (GO) enrichment analysis results for each of the nine protein expression trajectories.

**Table S5. Isoform Switching Dynamics and Functional Consequences. Related to** Figure 4. This table provides the complete isoform-level differential analysis results and the characterization of their functional consequences, as summarized in Figure 4.

● **DTE (Differential Transcript Expression):** Statistical results (log2FC, FDR) for all isoforms with significant changes in abundance (∼49,000 events).

● **DTU (Differential Transcript Usage):** Statistical results (change in isoform fraction ’dIF’, FDR) for all isoforms with significant changes in relative usage (∼24,000 events).

● **Summarized/Detailed switch conseq/AS:** Gene-level and isoform switch-level characterization of the alternative splicing (e.g., microexon inclusion, intron retention) and functional consequences (e.g., NMD status, protein domain gain/loss) of isoform switching.

● **SFARI Gene Enrichment:** Statistical results (odds ratio, *P*-value) for the enrichment of SFARI ASD risk genes among genes undergoing each class of switching event, as shown in Figure 4C.

● **GO Term Enrichment:** Complete Gene Ontology (GO) enrichment analysis results for genes regulated by specific switching events (e.g., intron retention), as shown in Figure 4D.

**Table S6. Coordination of Alternative Splicing (AS) and Alternative Polyadenylation (APA). Related to** Figure 5 **and Figure S8.** This table provides the complete statistical results and supporting data for the analysis of AS-APA coordination and 3’UTR dynamics.

● **Quasi - Results & EMMs:** Contains the full statistical output from the primary quasi-binomial generalized linear model (GLM) approach. This includes the 2,789 significant coordination events, their FDR values for ’global’ and ’interaction’ effects, and the estimated marginal means (EMMs) values.

● **Fishers - Results:** Contains the complete results from the validation analysis using the contingency table-based Fisher’s exact test method, for comparison.

● **QAPA & APA:** Includes the output from the QAPA analysis and the custom long-read APA analysis, detailing proximal polyA site usage (PPAU) and the comparison of dPPAU to gene expression (log2FC) used to generate **Figure S8**.

**Table S7. Burden of Disruptive *de novo* Mutations in ASD Cohorts. Related to** Figure 6. This table provides QC attributes related to all variants used for the *de novo* variant (DNV) burden analysis, as summarized in Figure 6B. The data is stratified by cohort (MSSNG, SSC, SPARK_WGS, and SPARK_WES) in the “data_cohort” column, and by variant category in the “is_known” column, which is a column that is “True” when the isoform is known, “False” when the isoform is novel.

● **Known:** Variants classified as disruptive using the GENCODE V47 reference.

● **Novel:** Variants reclassified as disruptive *only* by this study’s novel transcriptome. This data is used to calculate the odds ratios (OR) and *P*-values (Fisher’s exact test) to assess the contribution of novel elements to ASD risk.

## Notes

### Competing Interest Statement

The authors have declared no competing interest.

## References

1. Furlanis, E. & Scheiffele, P. Regulation of Neuronal Differentiation, Function, and Plasticity by Alternative Splicing. Annu. Rev. Cell Dev. Biol. 34, 451–469 (2018).

2. Gandal, M. J. et al. Transcriptome-wide isoform-level dysregulation in ASD, schizophrenia, and bipolar disorder. Science 362, (2018).

3. Irimia, M. et al. A Highly Conserved Program of Neuronal Microexons Is Misregulated in Autistic Brains. Cell 159, 1511–1523 (2014).

4. Parikshak, N. N. et al. Genome-wide changes in lncRNA, splicing, and regional gene expression patterns in autism. Nature 540, 423–427 (2016).

5. Cummings, B. B. et al. Improving genetic diagnosis in Mendelian disease with transcriptome sequencing. Sci. Transl. Med. 9, eaal5209 (2017).

6. Nagura, Y. et al. Long-read sequencing reveals novel isoform-specific eQTLs and regulatory mechanisms of isoform expression in human B cells. Genome Biol. 26, 110 (2025).

7. Stark, R., Grzelak, M. & Hadfield, J. RNA sequencing: the teenage years. Nat. Rev. Genet. 20, 631–656 (2019).

8. Steijger, T. et al. Assessment of transcript reconstruction methods for RNA-seq. Nat. Methods 10, 1177–1184 (2013).

9. Pardo-Palacios, F. J. et al. SQANTI3: curation of long-read transcriptomes for accurate identification of known and novel isoforms. Nat. Methods 1–5 (2024) doi:10.1038/s41592-024-02229-2.

10. Hardwick, S. A. et al. Single-nuclei isoform RNA sequencing unlocks barcoded exon connectivity in frozen brain tissue. Nat. Biotechnol. 40, 1082–1092 (2022).

11. Zhang, Z., Bae, B., Cuddleston, W. H. & Miura, P. Coordination of alternative splicing and alternative polyadenylation revealed by targeted long read sequencing. Nat. Commun. 14, 5506 (2023).

12. Bamford, R. A. et al. An atlas of expressed transcripts in the prenatal and postnatal human cortex. 2024.05.24.595768 Preprint at 10.1101/2024.05.24.595768 (2024).

13. Patowary, A. et al. Developmental isoform diversity in the human neocortex informs neuropsychiatric risk mechanisms. Science 384, eadh7688 (2024).

14. Xu, J. et al. Long-read RNA-sequencing reveals transcript-specific regulation in human-derived cortical neurons. Open Biol. 15, 250200 (2025).

15. Al’Khafaji, A. M. et al. High-throughput RNA isoform sequencing using programmed cDNA concatenation. Nat. Biotechnol. 1–5 (2023) doi:10.1038/s41587-023-01815-7.

16. Wissel, D. et al. A Systematic Benchmark of High-Accuracy PacBio Long-Read RNA Sequencing for Transcript-Level Quantification.

17. Fernando, M. B., Ahfeldt, T. & Brennand, K. J. Modeling the complex genetic architectures of brain disease. Nat. Genet. 52, 363–369 (2020).

18. Lin, H. C. et al. NGN2 induces diverse neuron types from human pluripotency. Stem Cell Rep. 16, 2118–2127 (2021).

19. Mudge, J. M. et al. GENCODE 2025: reference gene annotation for human and mouse. Nucleic Acids Res. 53, D966–D975 (2025).

20. Tardaguila, M. et al. SQANTI: extensive characterization of long-read transcript sequences for quality control in full-length transcriptome identification and quantification. Genome Res. 28, 396–411 (2018).

21. Tapial, J. et al. An atlas of alternative splicing profiles and functional associations reveals new regulatory programs and genes that simultaneously express multiple major isoforms. Genome Res. 27, 1759–1768 (2017).

22. Varabyou, A., Erdogdu, B., Salzberg, S. L. & Pertea, M. Investigating open reading frames in known and novel transcripts using ORFanage. Nat. Comput. Sci. 3, 700–708 (2023).

23. Miller, R. M. et al. Enhanced protein isoform characterization through long-read proteogenomics. Genome Biol. 23, 69 (2022).

24. Humphrey, J. et al. Long-read RNA sequencing atlas of human microglia isoforms elucidates disease-associated genetic regulation of splicing. Nat. Genet. 1–12 (2025) doi:10.1038/s41588-025-02099-0.

25. Duffy, E. E. et al. Developmental dynamics of RNA translation in the human brain. Nat. Neurosci. 25, 1353–1365 (2022).

26. Hulme, A. J., Maksour, S., Glover, M. S.-C., Miellet, S. & Dottori, M. Making neurons, made easy: The use of Neurogenin-2 in neuronal differentiation. Stem Cell Rep. 17, 14–34 (2022).

27. Vitting-Seerup, K. & Sandelin, A. The Landscape of Isoform Switches in Human Cancers. Mol. Cancer Res. 15, 1206–1220 (2017).

28. Blair, J. D., Hockemeyer, D., Doudna, J. A., Bateup, H. S. & Floor, S. N. Widespread Translational Remodeling during Human Neuronal Differentiation. Cell Rep. 21, 2005–2016 (2017).

29. Kwak, P. B. & Tomari, Y. The N domain of Argonaute drives duplex unwinding during RISC assembly. Nat. Struct. Mol. Biol. 19, 145–151 (2012).

30. Zhang, X. et al. Cell-Type-Specific Alternative Splicing Governs Cell Fate in the Developing Cerebral Cortex. Cell 166, 1147–1162.e15 (2016).

31. Tian, B. & Manley, J. L. Alternative polyadenylation of mRNA precursors. Nat. Rev. Mol. Cell Biol. 18, 18–30 (2017).

32. Anvar, S. Y. et al. Full-length mRNA sequencing uncovers a widespread coupling between transcription initiation and mRNA processing. Genome Biol. 19, 46 (2018).

33. Spies, N., Burge, C. B. & Bartel, D. P. 3′ UTR-isoform choice has limited influence on the stability and translational efficiency of most mRNAs in mouse fibroblasts. Genome Res. 23, 2078–2090 (2013).

34. Ha, K. C. H., Blencowe, B. J. & Morris, Q. QAPA: a new method for the systematic analysis of alternative polyadenylation from RNA-seq data. Genome Biol. 19, 45 (2018).

35. Kiltschewskij, D. J., Harrison, P. F., Fitzsimmons, C., Beilharz, T. H. & Cairns, M. J. Extension of mRNA poly(A) tails and 3′UTRs during neuronal differentiation exhibits variable association with post-transcriptional dynamics. Nucleic Acids Res. 51, 8181–8198 (2023).

36. Lord, J. et al. Pathogenicity and selective constraint on variation near splice sites. Genome Res. 29, 159–170 (2019).

37. Gudmundsson, S. et al. Exploring penetrance of clinically relevant variants in over 800,000 humans from the Genome Aggregation Database. Nat. Commun. 16, 9623 (2025).

38. Karczewski, K. J. et al. The mutational constraint spectrum quantified from variation in 141,456 humans. Nature 581, 434–443 (2020).

39. Feliciano, P. et al. Exome sequencing of 457 autism families recruited online provides evidence for autism risk genes. Npj Genomic Med. 4, 19 (2019).

40. Fischbach, G. D. & Lord, C. The Simons Simplex Collection: A Resource for Identification of Autism Genetic Risk Factors. Neuron 68, 192–195 (2010).

41. Trost, B. et al. Genomic architecture of autism from comprehensive whole-genome sequence annotation. Cell 185, 4409–4427.e18 (2022).

42. Barbosa-Morais, N. L. et al. The Evolutionary Landscape of Alternative Splicing in Vertebrate Species. Science 338, 1587–1593 (2012).

43. Lek, M. et al. Analysis of protein-coding genetic variation in 60,706 humans. Nature 536, 285–291 (2016).

44. Tress, M. L., Abascal, F. & Valencia, A. Alternative Splicing May Not Be the Key to Proteome Complexity. Trends Biochem. Sci. 42, 98–110 (2017).

45. Raj, A. & van Oudenaarden, A. Nature, nurture, or chance: stochastic gene expression and its consequences. Cell 135, 216–226 (2008).

46. Wan, Y. & Larson, D. R. Splicing heterogeneity: separating signal from noise. Genome Biol. 19, 86 (2018).

47. Dai, A., Temporal, S. & Schulz, D. J. Cell-specific patterns of alternative splicing of voltage-gated ion channels in single identified neurons. Neuroscience 168, 118–129 (2010).

48. Lambourne, L. et al. Widespread variation in molecular interactions and regulatory properties among transcription factor isoforms. Mol. Cell 85, 1445–1466.e13 (2025).

49. Deshwar, A. R. et al. Trio RNA sequencing in a cohort of medically complex children. Am. J. Hum. Genet. 110, 895–900 (2023).

50. Stark, J. C. et al. Clinical applications of and molecular insights from RNA sequencing in a rare disease cohort. Genome Med. 17, 72 (2025).

51. Jaramillo Oquendo, C., et al. Identification of diagnostic candidates in Mendelian disorders using an RNA sequencing-centric approach. Genome Med. 16, 110 (2024).

52. Clavell-Revelles, P. et al. Long-read transcriptomics of a diverse human cohort reveals widespread ancestry bias in gene annotation. Preprint at 10.1101/2025.03.14.643250 (2025).

53. Hildebrandt, M. R. et al. Precision Health Resource of Control iPSC Lines for Versatile Multilineage Differentiation. Stem Cell Rep. 13, 1126–1141 (2019).

54. Tian, R. et al. CRISPR Interference-Based Platform for Multimodal Genetic Screens in Human iPSC-Derived Neurons. Neuron 104, 239–255.e12 (2019).

55. Li, Y. et al. Induction of Expansion and Folding in Human Cerebral Organoids. Cell Stem Cell 20, 385–396.e3 (2017).

56. Ran, F. A. et al. Genome engineering using the CRISPR-Cas9 system. Nat. Protoc. 8, 2281–2308 (2013).

57. Autar, K. et al. A functional hiPSC-cortical neuron differentiation and maturation model and its application to neurological disorders. Stem Cell Rep. 17, 96–109 (2021).

58. Nehme, R. et al. Combining NGN2 Programming with Developmental Patterning Generates Human Excitatory Neurons with NMDAR-Mediated Synaptic Transmission. Cell Rep. 23, 2509–2523 (2018).

59. Danecek, P. et al. Twelve years of SAMtools and BCFtools. GigaScience 10, giab008 (2021).

60. Abugessaisa, I. et al. refTSS: A Reference Data Set for Human and Mouse Transcription Start Sites. J. Mol. Biol. 431, 2407–2422 (2019).

61. Herrmann, C. J. et al. PolyASite 2.0: a consolidated atlas of polyadenylation sites from 3′ end sequencing. Nucleic Acids Res. 48, D174–D179 (2020).

62. Dobin, A. et al. STAR: ultrafast universal RNA-seq aligner. Bioinformatics 29, 15–21 (2013).

63. The UniProt Consortium. UniProt: the Universal Protein Knowledgebase in 2023. Nucleic Acids Res. 51, D523–D531 (2023).

64. Eng, J. K. et al. A Deeper Look into Comet—Implementation and Features. J. Am. Soc. Mass Spectrom. 26, 1865–1874 (2015).

65. Käll, L., Canterbury, J. D., Weston, J., Noble, W. S. & MacCoss, M. J. Semi-supervised learning for peptide identification from shotgun proteomics datasets. Nat. Methods 4, 923–925 (2007).

66. Käll, L., Storey, J. D., MacCoss, M. J. & Noble, W. S. Assigning Significance to Peptides Identified by Tandem Mass Spectrometry Using Decoy Databases. J. Proteome Res. 7, 29–34 (2008).

67. Brademan, D. R., Riley, N. M., Kwiecien, N. W. & Coon, J. J. Interactive Peptide Spectral Annotator: A Versatile Web-based Tool for Proteomic Applications. Mol. Cell. Proteomics MCP 18, S193–S201 (2019).

68. Brar, G. A. & Weissman, J. S. Ribosome profiling reveals the what, when, where and how of protein synthesis. Nat. Rev. Mol. Cell Biol. 16, 651–664 (2015).

69. Kechin, A., Boyarskikh, U., Kel, A. & Filipenko, M. cutPrimers: A New Tool for Accurate Cutting of Primers from Reads of Targeted Next Generation Sequencing. J. Comput. Biol. J. Comput. Mol. Cell Biol. 24, 1138–1143 (2017).

70. Anders, S., Pyl, P. T. & Huber, W. HTSeq--a Python framework to work with high-throughput sequencing data. Bioinforma. Oxf. Engl. 31, 166–169 (2015).

71. Varabyou, A., Erdogdu, B., Salzberg, S. L. & Pertea, M. Investigating open reading frames in known and novel transcripts using ORFanage. Nat. Comput. Sci. 3, 700–708 (2023).

72. Chen, Y., Chen, L., Lun, A. T. L., Baldoni, P. L. & Smyth, G. K. edgeR v4: powerful differential analysis of sequencing data with expanded functionality and improved support for small counts and larger datasets. Nucleic Acids Res. 53, (2025).

73. Love, M. I., Huber, W. & Anders, S. Moderated estimation of fold change and dispersion for RNA-seq data with DESeq2. Genome Biol. 15, 550 (2014).

74. Vitting-Seerup, K. & Sandelin, A. IsoformSwitchAnalyzeR: analysis of changes in genome-wide patterns of alternative splicing and its functional consequences. Bioinformatics 35, 4469–4471 (2019).

75. Vitting-Seerup, K., Porse, B. T., Sandelin, A. & Waage, J. spliceR: an R package for classification of alternative splicing and prediction of coding potential from RNA-seq data. BMC Bioinformatics 15, 81 (2014).

76. Punta, M. et al. The Pfam protein families database. Nucleic Acids Res. 40, D290–D301 (2012).

77. Mészáros, B., Erdős, G. & Dosztányi, Z. IUPred2A: context-dependent prediction of protein disorder as a function of redox state and protein binding. Nucleic Acids Res. 46, W329–W337 (2018).

78. Huang, T. et al. MSstatsTMT: Statistical Detection of Differentially Abundant Proteins in Experiments with Isobaric Labeling and Multiple Mixtures. Mol. Cell. Proteomics 19, 1706–1723 (2020).

79. Consortium, G. O. et al. The Gene Ontology knowledgebase in 2023. Genetics 224, iyad031 (2023).

80. Kolberg, L., Raudvere, U., Kuzmin, I., Vilo, J. & Peterson, H. gprofiler2 – an R package for gene list functional enrichment analysis and namespace conversion toolset g:Profiler. F1000Research 9, 709 (2020).

81. Trincado, J. L. et al. SUPPA2: fast, accurate, and uncertainty-aware differential splicing analysis across multiple conditions. Genome Biol. 19, 40 (2018).

82. Lenth, R. V. Emmeans: Estimated Marginal Means, Aka Least-Squares Means. (2025).

83. McLaren, W. et al. The Ensembl Variant Effect Predictor. Genome Biol. 17, 122 (2016).

84. Wang, K., Li, M. & Hakonarson, H. ANNOVAR: functional annotation of genetic variants from high-throughput sequencing data. Nucleic Acids Res. 38, e164–e164 (2010).

85. Blakes, A. J. M. et al. A systematic analysis of splicing variants identifies new diagnoses in the 100,000 Genomes Project. Genome Med. 14, 79 (2022).

86. Samocha, K. E. et al. A framework for the interpretation of de novo mutation in human disease. Nat. Genet. 46, 944–950 (2014).

